# Childhood Maltreatment and Deviations from Normative Brain Structure: Results from 3,711 Individuals from the ENIGMA MDD and ENIGMA PTSD

**DOI:** 10.1101/2025.06.04.657717

**Authors:** Tiffany C. Ho, the ENIGMA MDD Working Group, ENIGMA PTSD Working Group

**Author notes:** Corresponding author Tiffany C. Ho, PhD Assistant Professor Department of Psychology Brain Research Institute Interdepartmental Graduate Program in Neuroscience University of California, Los Angeles A326B Psychology Building, Los Angeles, CA, 90095. A list of working groups authors and their affiliations appears at the end of the paper.

## Abstract

Childhood maltreatment (CM), encompassing abuse and neglect, affects over two-thirds of the general population and increases risk for stress-related psychopathology, including major depressive disorder (MDD) and posttraumatic stress disorder (PTSD). The extent to which neuroanatomical alterations in MDD and PTSD are attributable to CM, however, is uncertain. Here, we analyzed CM and 3D structural brain MRI data from 3,711 participants in the ENIGMA MDD and PTSD Working Groups (25 sites; 33.3±13.0 years; 59.9% female). Normative modeling estimated deviation z-scores for 14 subcortical volumes (SV), 68 cortical thickness (CT), and 68 surface area (SA) measures, capturing differences from population norms. Transdiagnostic associations between CM and brain deviation scores were evaluated within each sex and age cohort. In young adults (ages 18-35), abuse was associated with larger volumes in thalamus and pallidum, thinner isthmus cingulate and middle frontal regions, and thicker medial orbitofrontal cortex; there were no significant effects in pediatric (≤18 years) participants. The strongest effects were observed in young female adults (|β|=.07-.22, *q*<.05): greater abuse and neglect were correlated with smaller hippocampus and putamen volumes, thinner entorhinal cortex, and smaller SA in fusiform/inferior parietal regions, and with larger SA in orbitofrontal and occipital cortices. In males, abuse had widespread effects on CT and SA (|β|=.1-.18, *q*<.05); effects for neglect were minimal. Our findings of age- and sex-specific instantiations of CM on brain morphometry highlight the importance of developmental context in understanding how adverse experiences shape neurobiological vulnerability to MDD and PTSD.

## INTRODUCTION

Early life adversity (ELA), particularly childhood maltreatment (CM) such as abuse and neglect, is common, affecting more than two-thirds of the general population [1, 2]. Although not all individuals who experience CM develop mental disorders, experiences of CM remains one of the most potent risk factors for stress-related psychopathologies, including major depressive disorder (MDD) and posttraumatic stress disorder (PTSD) [3–5]. Importantly, individuals who are diagnosed with such disorders and have a history of CM tend to experience an earlier age of onset, poorer clinical outcomes, and greater treatment resistance than those with these disorders but no history of CM [6]. Despite the clear link between CM experience and subsequent risk for psychiatric disorders, the neurobiological mechanisms underlying this association are not well understood. This knowledge gap is notable given regional brain differences observed in deviations from normative neurodevelopment and the lack of studies adopting a lifespan approach [7].

Specifically, many studies examining the effects of CM on brain morphometry have not adequately accounted for the potential influence of CM on inter-individual variation in brain morphometry across regions [8]. Brain regions exhibit diverse developmental courses, with some following inverted U-shaped trajectories that reach their peak in childhood or adolescence, with evidence of sex differences in these trajectories [9–12]. Consequently, interpreting adversity-related effects necessitates consideration of inter-individual variation specific to age and sex [13, 14]. Two example regions are thalamus and amygdala, which are consistently implicated in stress and trauma-related processes and show robust associations with CM in previous research [15, 16]. Larger thalamic volumes during childhood may suggest accelerated maturation, but the same pattern observed in adulthood may reflect delayed or atypical development [9, 17]. Additionally, while both male and female brains show inverted U-shaped age-related changes in amygdala volume, males demonstrate a more rapid increase in amygdala size during childhood compared to females, leading to sexual dimorphism and inter-individual variation. To advance our understanding of inter-individual variation in brain morphometry, investigators have recently charted normative brain growth, akin to pediatric growth charts, to quantify individual variation in brain morphometry with respect to age stratified normative data [10, 18]. Such an approach allows for a more nuanced understanding of how CM may influence inter-individual variation in brain morphometric development [19].

In addition, the effect of CM exposure on brain phenotypes may depend on developmental stage [20]. Germane to this idea is the concept of *sensitive periods*, which are periods of heightened plasticity during which environmental influences have outsized effects on developing systems [7]. Rather than standing in strict opposition to an accumulation model—which posits that effects scale with the number of exposures that typically increase with age [21]—these perspectives may be complementary, as adversity experienced during a sensitive window could confer a large initial impact that is compounded later in life with further exposures. Indeed, childhood maltreatment has been associated with an increased risk of psychiatric disorders that persist well into adulthood. To date, however, few studies have addressed this issue using samples from across the lifespan. Within a sensitive period, the neurobiological sequelae of adverse experiences may be most prominent immediately after exposure (as described in *recency* models [22]), or they may not be manifested or measurable at the phenotypic level until later in development [23] (as described in *incubation* models [24, 25]). For example, several empirical studies have documented this delayed manifestation pattern, finding minimal structural brain differences in children exposed to ELA, but more pronounced effects emerging in adolescence and early adulthood [20, 26]. To distinguish among these possibilities, a lifespan approach is critically needed [21].

Finally, prior research on the structural brain correlates of CM has not fully captured the nuanced effects of different experience types [27, 28] or potential sex differences in these associations [29]. Recent data-driven approaches have identified distinct co-occurring dimensions of ELA that may have differential neural correlates [30], though these dimensional patterns have not yet been examined in relation to normative brain development. Prominent theoretical frameworks distinguish between adversity characterized by threat versus deprivation and also highlight the impact of fragmented care—defined as inconsistent, unpredictable, or disrupted caregiving—on neurodevelopment [31]. Various forms of childhood maltreatment—such as emotional abuse, emotional neglect, physical abuse, physical neglect, and sexual abuse—may lead to distinct cognitive and behavioral adaptations or maladaptive coping strategies [32, 33]. For example, sexual abuse, which typically involves high levels of threat, has been associated with structural deficits in reward circuits and altered amygdala reactivity [16, 34, 35], whereas emotional maltreatment, often linked to deprivation, has been connected with abnormalities in fronto-limbic socioemotional networks [36]. However, the current evidence remains incomplete.

The impact of childhood maltreatment likely interacts with hormonal factors and gender-specific social environments, potentially resulting in differing outcomes for males and females. For example, women are more than twice as likely as men to develop PTSD following exposures to all types of abuse and neglect [37, 38]; similar sex differences have been observed in depression, with women showing higher prevalence rates [39, 40]. Despite this, many studies have relied on mixed-sex samples, often due to limited sample sizes, missing opportunities to detect sex differences in the neurobiological sequelae of ELA.

Taken together, there is a clear need to examine the enduring effects of CM on brain morphometry across the lifespan. In this study, we employed a normative modeling approach that quantifies individual deviations from expected neuroanatomical variation by generating continuous z-scores, which indicate the direction of normative deviation (with lower scores reflecting values below the normative range, and higher scores reflecting values above the normative range) [41, 42]. Normative modeling uniquely addresses methodological challenges of accounting for developmental heterogeneity by providing age- and sex-adjusted reference standards analogous to pediatric growth charts, thus offering advantages over traditional approaches that rely on group-average comparisons. Our aim was to explore the extent to which the pattern of brain deviations is correlated with continuous measures of childhood trauma severity in a way that aligns with dimensional models of ELA. In our framework, different types of maltreatment—such as abuse and neglect—are expected to differentially impact distinct neural circuits in parallel. For instance, whereas threat-related abuse may predominantly affect regions involved in stress regulation and fear learning (e.g., the hippocampus and amygdala), deprivation-related neglect may influence circuits underpinning reward processing and socioemotional function (e.g., the striatum, medial orbitofrontal cortex, and association cortices). Recognizing that brain development and the neurobiological sequelae of CM vary by age and sex, we analyzed cortical thickness, surface area, and subcortical volumes separately for different age cohorts and for males and females. Our large, trauma- enriched sample (N = 3,711), spanning ages 8 to 69 from 25 international sites via the ENIGMA MDD and PTSD Working Groups [43–45], allowed us to correlate continuous measures of trauma severity with these continuous deviation scores. This approach aims to determine the extent to which our findings support a dimensional model of CM that captures differential neural alterations associated with different forms of adversity, and how these associations vary as a function of age and sex.

## METHODS ENIGMA Data

Data from 25 independent sites (Fig. S1) across eight countries were obtained through the ENIGMA Consortium Major Depressive Disorder (MDD) and Posttraumatic Stress Disorder (PTSD) Working Groups. The overall sample consisted of 1,389 patients (872 MDD, 517 PTSD), and 2,322 healthy controls. Cortical thickness and cortical surface area measures were extracted based on the Desikan-Killiany (aparc) atlas [46], while subcortical volumes were segmented using the Aseg atlas in FreeSurfer [47]. All neuroimaging data were processed with FreeSurfer according to the standardized processing pipelines and quality assurance procedures stipulated by the ENIGMA Consortium (https://enigma.ini.usc.edu/protocols/imaging-protocols/). Detailed information on participant recruitment, data collection procedures, and MRI acquisition and preprocessing for each site is provided in Supplementary Methods S1-S2.

### Childhood Maltreatment Assessment

Childhood Maltreatment (CM) was assessed in all samples using the Childhood Trauma Questionnaire-Short Form (CTQ-SF) [48], a widely validated retrospective self-report measure of childhood maltreatment. The CTQ-SF comprises five subscales: physical abuse, emotional abuse, sexual abuse, physical neglect, and emotional neglect. Two composite scores were derived: (1) total severity of childhood abuse, calculated as the sum of the three abuse subscales, and (2) total severity of childhood neglect, calculated as the sum of the two neglect subscales. We separated abuse from neglect to facilitate the detection of distinct neurobiological correlates associated with each type of maltreatment.

### Normative Modeling of Brain Morphometry

Sex-specific normative models for each FreeSurfer-derived regional subcortical volume, cortical thickness, and surface area measure were generated using the CentileBrain normative modeling tools (Supplementary Methods S3) developed by the ENIGMA Lifespan Group [49] and are freely available (https://centilebrain.org/). These models were developed using data from an independent multi-site sample of 37,407 healthy individuals (53.3% female; ages 3-90 years and provide separate standards for females and males. CentileBrain generated z-scores for each morphometric measure in each participant, reflecting deviations from age- and sex-specific population means. Z-scores were calculated by subtracting the brain measure estimated by CentileBrain from the raw value, then dividing by the model’s root mean square error. While the CentileBrain algorithms include site harmonization based on Combat-GAM [50], we did not use this feature for reasons discussed in the Sensitivity Analysis *Harmonization Effects* section.

### Statistical Analyses

*Main Analyses.* We pooled the extracted subcortical volumes, cortical thickness, and cortical surface area measures from individual subjects across all sites from the ENIGMA MDD and PTSD Working Groups into a single database. General linear models examined associations between CTQ scores (total abuse and neglect) and z-scores for each regional morphometric measure (14 subcortical volumes, 68 cortical thickness, 68 surface area measures; separately for left and right hemispheres). Separate analyses were conducted for males and females. As the z-scores already account for age and intracranial volume in the normative model, we did not further adjust for these variables. CTQ scores were not associated with age, age-squared, or sex.

*Age-Dependent Effects*. While normative modeling inherently accounts for age by generating deviation scores relative to a population-derived reference for each specific age (akin to pediatric growth charts), we stratified our sample into pediatric (≤18 years, n=349), young adult (>18 and <35 years, n=1,908), and older adult (≥35 years, n=1,454) cohorts to explicitly examine developmental stages that correspond to distinct phases of neurodevelopment and plasticity. Specifically, categorizing participants into these broadly defined age cohorts allows us to investigate potential differences in the magnitude and specificity of ELA-associated brain deviations during critical developmental windows, such as adolescence versus early and later adulthood. Moreover, this stratification approach is consistent with methods used in several prior ENIGMA Consortium studies [51–54], allowing for comparability across multiple large-scale neuroimaging investigations.

*Multiple Comparisons Correction.* To adjust for multiple comparisons, we applied a false discovery rate (FDR) correction using the Benjamini-Hochberg procedure, with *p*-values adjusted separately for each age cohort and sex and for each metric (subcortical volumes, cortical thickness, cortical surface area).

Results were considered significant if the FDR-corrected *p*-value (*q*) ≤0.05. The number of comparisons was 68 for cortical thickness and surface area analyses, and 14 for subcortical volume analyses. To obtain comparable standardized regression coefficients, z-scores for each region served as outcome measures.

*Sex and Age Differences in Brain-CM Associations.* To statistically compare the strength of associations between sexes and age cohorts, we employed Fisher’s exact transformation tests (Supplementary Methods S4). This method allowed us to determine whether the standardized regression coefficients (β) from our general linear models differed significantly between males and females (within the same age cohort) and between young adults and older adults (within the same sex).

### Sensitivity Analysis

*Diagnostic Effects.* We adjusted for diagnostic status ("patients" versus "controls") as a covariate in our models. This binary grouping was preferred over separating MDD and PTSD due to recruitment variability across sites and potential symptom overlap. For more details, see Supplementary Methods S5.

*Site-Specific Effects.* Leave-one-out cross-validation assessed site-specific influences by sequentially excluding each site and re-running our models. We used the ranges of significance values (*p*-values) to evaluate stability of results across sites. For more details, see Supplementary Methods S6.

*Harmonization Effects.* Although CentileBrain methods effectively addressed cross-site variation and eliminated the need for harmonization [10], out of precaution we tested the effect of harmonization on our dataset using ComBat-GAM (Supplementary Methods S7). The results were similar with the main results reported in this manuscript (Fig. S4). Some sites lacked a cohort of healthy controls, likely resulting in meaningful variance associated with CM and diagnosis being removed by ComBat-GAM due to conflation with site effects [50]. We therefore opted to report the non-harmonized results in the main manuscript.

*Power Analysis for Pediatric Cohort.* Bootstrap sampling analysis determined whether null findings in the pediatric cohort stemmed from limited statistical power rather than developmental differences (Supplementary Methods S8). Power was calculated as the proportion of bootstrap iterations yielding significant results (*p*<.05).

## RESULTS

### Participant Characteristics and Childhood Trauma

The final sample demographics and clinical characteristics, stratified by age cohorts, are presented in Table S1. Sample measures are further broken down by diagnostic group, age, and sex respectively (Tables S2-S3, Fig. S1-3).

### Associations Between CM and Subcortical Volume Deviation Scores

Our analyses revealed significant associations between CM and deviations from normative subcortical volume (SV) in adults, with distinct patterns observed across age cohorts, sexes, and CM types (Fig. 1; Table 1). Notably, there were no significant associations between CM and deviations in normative SV in the pediatric cohort.

**Figure 1.**
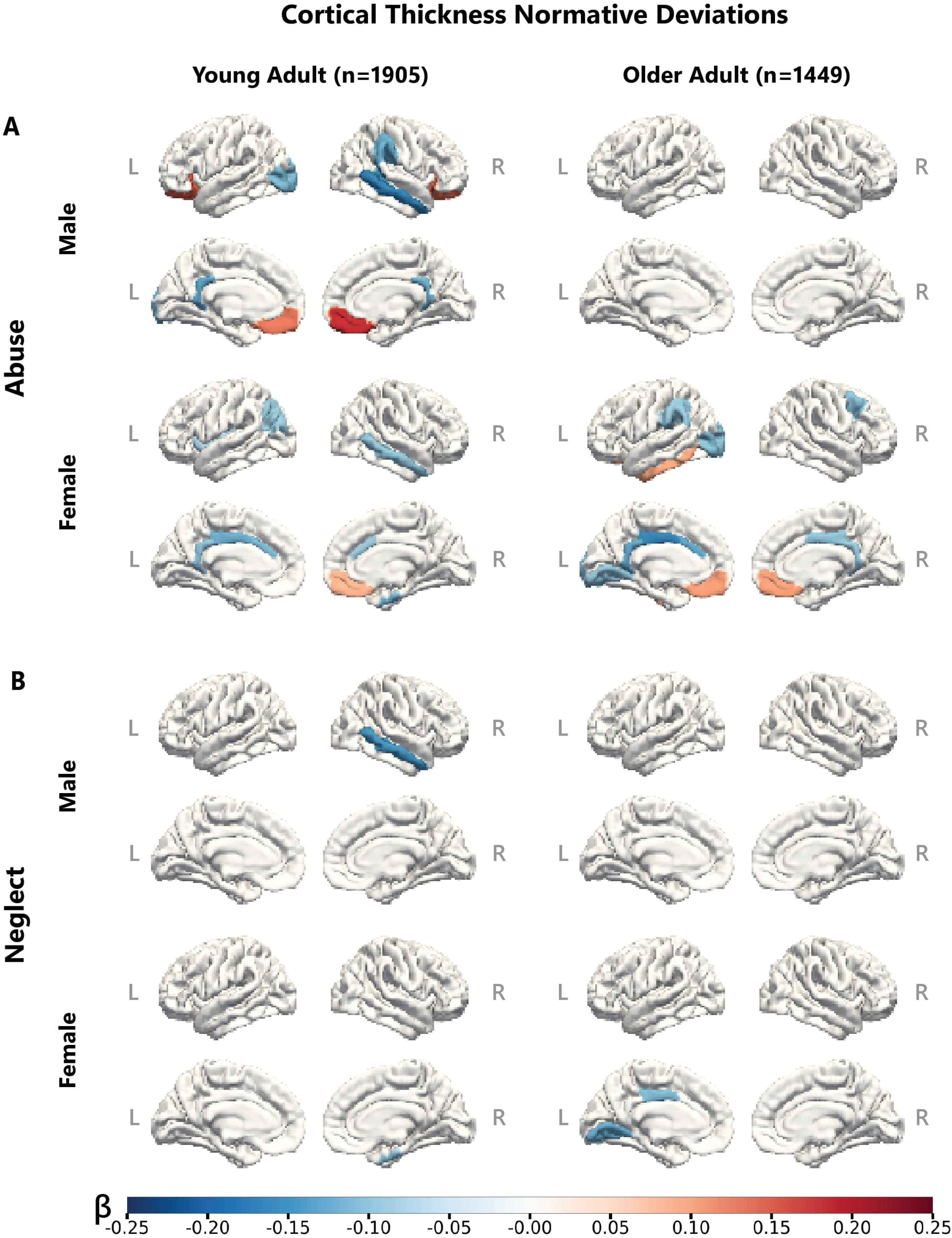
Deviations from Normative Subcortical Volume Associated with Childhood Trauma. **A.** Normative deviations in subcortical volumes associated with abuse in males and females; **B.** Normative deviations in subcortical volumes associated with neglect in males and females. β represents the standardized effect size from the general linear model (GLM, FDR corrected *q*<.05). Results from the pediatric cohort are not displayed as there were no significant associations between CTQ and subcortical volumes in this age cohort.

**Table 1.** Deviations from Normative Subcortical Volume Associated with Childhood Trauma. P- values are **t**hresholded to show regions with normative values significantly associated with childhood trauma in at least one age cohort. See the full results table of all regions and cohorts in Table S4. Significant regions (*q*<.05) are indicated by bolded effect sizes and *p* values. β represents the standardized effect size from the GLM. Results from the pediatric cohort are not displayed as there were no significant associations between CTQ and subcortical volumes in this age cohort.

| Childhood Trauma | Sex | Region | Young Adult (n=1908) |  | Older Adult (n=1454) |  |
| --- | --- | --- | --- | --- | --- | --- |
| | | | $\beta$ (SE) | *p* value | $\beta$ (SE) | *p* value |
| Abuse | Male | Left Thalamus | **0.126 (0.036)** | **5.56e-04** | -0.026 (0.041) | 5.37e-01 |
|  |  | Right Thalamus | **0.1 (0.037)** | **6.32e-03** | -0.034 (0.041) | 4.16e-01 |
|  |  | Left Pallidum | **0.143 (0.036)** | **9.40e-05** | **0.117 (0.041)** | **4.51e-03** |
|  |  | Right Pallidum | **0.171 (0.036)** | **2.76e-06** | **0.114 (0.041)** | **5.74e-03** |
|  |  | Left Amygdala | -0.043 (0.037) | 2.46e-01 | **-0.104 (0.041)** | **1.21e-02** |
|  |  | Right Amygdala | 0.007 (0.037) | 8.56e-01 | **-0.12 (0.041)** | **3.54e-03** |
|  | Female | Right Thalamus | **0.11 (0.029)** | **1.62e-04** | **0.141 (0.034)** | **3.01e-05** |
|  |  | Left Caudate | **-0.065 (0.029)** | **2.69e-02** | -0.07 (0.034) | 3.93e-02 |
|  |  | Left Putamen | **-0.139 (0.029)** | **2.05e-06** | **-0.155 (0.034)** | **4.27e-06** |
|  |  | Right Putamen | **-0.106 (0.029)** | **2.74e-04** | -0.05 (0.034) | 1.45e-01 |
|  |  | Left Pallidum | **0.164 (0.029)** | **1.79e-08** | **0.146 (0.034)** | **1.55e-05** |
|  |  | Right Pallidum | **0.169 (0.029)** | **6.17e-09** | **0.115 (0.034)** | **7.29e-04** |
|  |  | Left Hippocampus | **-0.08 (0.029)** | **6.53e-03** | -0.019 (0.034) | 5.68e-01 |
|  |  | Right Hippocampus | **-0.129 (0.029)** | **1.09e-05** | -0.038 (0.034) | 2.69e-01 |
| Neglect | Male | - | - | - | - | - |
|  | Female | Right Thalamus | 0.064 (0.029) | 2.96e-02 | **0.096 (0.034)** | **4.64e-03** |
|  |  | Left Caudate | -0.032 (0.029) | 2.71e-01 | **-0.087 (0.034)** | **1.06e-02** |
|  |  | Right Caudate | -0.039 (0.029) | 1.78e-01 | **-0.086 (0.034)** | **1.08e-02** |
|  |  | Left Putamen | **-0.073 (0.029)** | **1.28e-02** | **-0.102 (0.034)** | **2.55e-03** |
|  |  | Left Pallidum | **0.114 (0.029)** | **9.09e-05** | 0.057 (0.034) | 9.41e-02 |
|  |  | Right Pallidum | **0.124 (0.029)** | **2.22e-05** | 0.032 (0.034) | 3.43e-01 |
|  |  | Right Hippocampus | **-0.092 (0.029)** | **1.68e-03** | 0.013 (0.034) | 6.98e-01 |

*Childhood Abuse.* In young adult males, more severe childhood abuse was associated with *larger* volumes in the bilateral thalamus and pallidum, which partially persisted in older adult males. Older adult males with more severe experiences of abuse exhibited *smaller* amygdala volumes. In young adult females, more severe childhood abuse was associated with *larger* volumes in the bilateral thalamus and pallidum; as well as *smaller* volumes of the bilateral hippocampus and putamen. This pattern persisted primarily in the right-hemisphere structures in the older adult females (Fig. 1A; Table 1).

*Childhood Neglect.* No significant associations were observed between childhood neglect and SV deviations in males across all age cohorts. In contrast, young adult females showed positive associations between more severe childhood neglect and *larger* bilateral pallidum volumes. Consistent with our analyses with childhood abuse, greater severity of neglect in young adult females was also associated with *smaller* volumes in the right hippocampus and left putamen. In older adult females, greater severity of neglect was associated with *smaller* volumes in the left putamen, *smaller* bilateral caudate volumes, and *larger* right thalamic volumes (Fig. 1B; Table 1).

### Associations Between Cortical Thickness Deviation Scores and ELA

Our analyses revealed significant associations between CM and deviations from normative cortical thickness (CT) in adults, with distinct patterns observed across age cohorts, sexes, and CM types (Fig. 2; Table 2). No significant associations were observed between CM and deviations from normative CT in the pediatric cohort.

**Figure 2.**
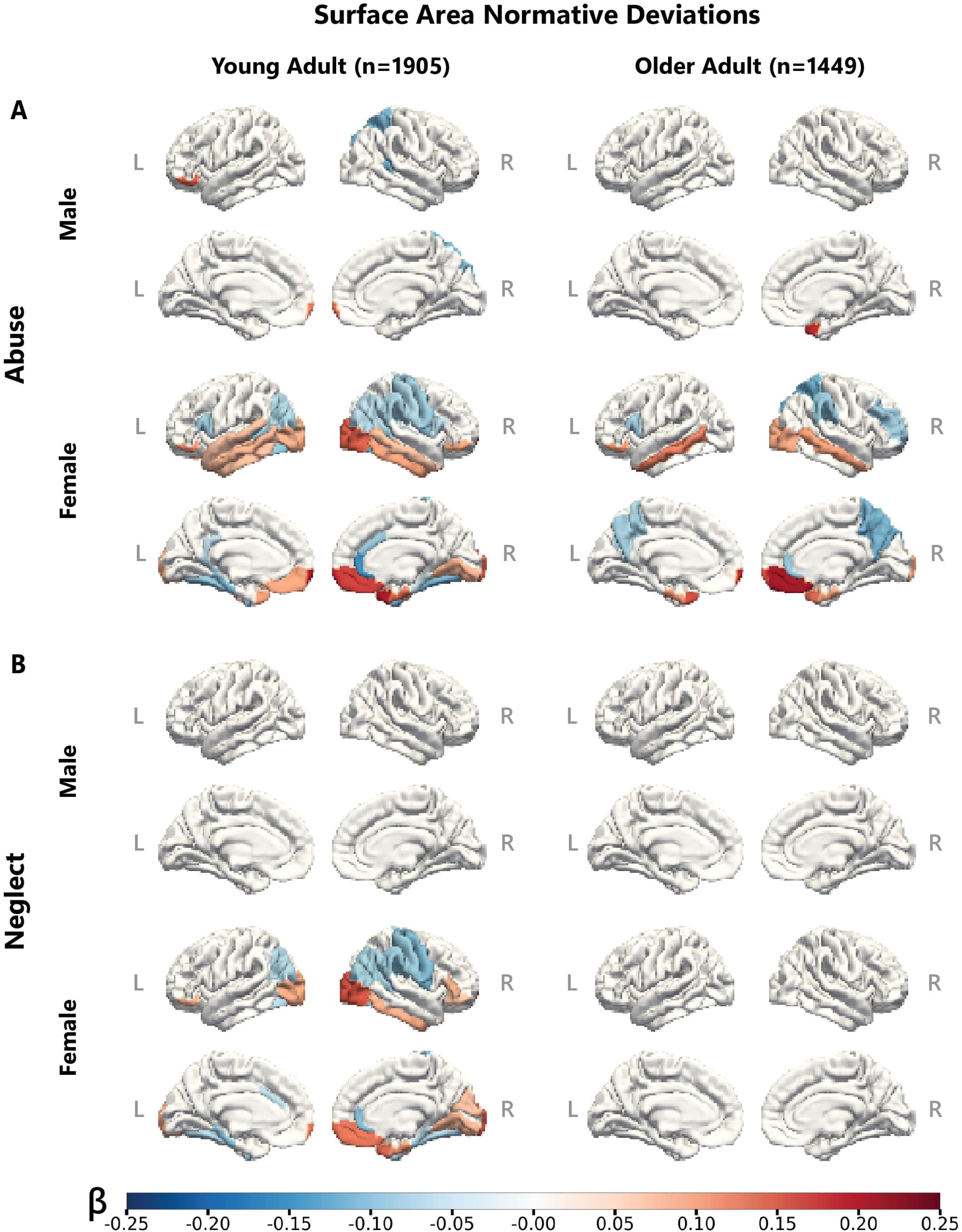
Deviations from Normative Cortical Thickness Associated with Childhood Trauma. **A.** Normative deviations in cortical thickness associated with abuse in males and females; **B.** Normative deviations in cortical thickness associated with neglect in males and females. β represents the standardized effect size from the GLM (FDR corrected *q*<.05). Results from the pediatric cohort are not displayed as there were no significant associations between CTQ and subcortical volumes in this age cohort.

**Table 2.** Deviations from Normative Cortical Thickness Associated with Childhood Trauma. P- values are **t**hresholded to show regions with normative values significantly associated with childhood trauma in at least one age cohort. See the full results table of all regions and cohorts in Supplementary Table S4. Significant regions (*q*<.05) are indicated by bolded effect sizes and *p* values. β represents the standardized effect size from the GLM. Results from the pediatric cohort are not displayed as there were no significant associations between CTQ and subcortical volumes in this age cohort.

| Childhood Trauma | Sex | Region | Young Adult (n=1905) |  | Older Adult (n=1449) |  |
| --- | --- | --- | --- | --- | --- | --- |
| | | | $\beta$ (SE) | *p* value | $\beta$ (SE) | *p* value |
| Abuse | Male | Left Isthmus cingulate | **-0.13 (0.036)** | **3.68e-04** | -0.017 (0.041) | 6.83e-01 |
|  |  | Left Lateral occipital | **-0.114 (0.037)** | **1.94e-03** | -0.066 (0.041) | 1.10e-01 |
|  |  | Left Lateral orbitofrontal | **0.152 (0.036)** | **3.40e-05** | -0.016 (0.041) | 7.07e-01 |
|  |  | Left Medial orbitofrontal | **0.132 (0.036)** | **3.15e-04** | 0.107 (0.041) | 9.40e-03 |
|  |  | Right Bankssts | **-0.153 (0.036)** | **2.83e-05** | -0.117 (0.041) | 4.65e-03 |
|  |  | Right Isthmus cingulate | **-0.118 (0.037)** | **1.25e-03** | -0.029 (0.041) | 4.84e-01 |
|  |  | Right Lateral orbitofrontal | **0.142 (0.036)** | **1.01e-04** | 0.013 (0.041) | 7.47e-01 |
|  |  | Right Medial orbitofrontal | **0.184 (0.036)** | **4.78e-07** | 0.102 (0.041) | 1.36e-02 |
|  |  | Right Middle temporal | **-0.164 (0.036)** | **6.85e-06** | -0.016 (0.041) | 6.99e-01 |
|  |  | Right Supramarginal | **-0.123 (0.036)** | **7.56e-04** | -0.061 (0.041) | 1.44e-01 |
|  | Female | Left Caudal anterior cingulate | **-0.116 (0.029)** | **7.59e-05** | **-0.132 (0.034)** | **9.95e-05** |
|  |  | Left Inferior parietal | **-0.095 (0.029)** | **1.15e-03** | -0.075 (0.034) | 2.83e-02 |
|  |  | Left Inferior temporal | 0.005 (0.029) | 8.69e-01 | **0.098 (0.034)** | **3.99e-03** |
|  |  | Left Isthmus cingulate | **-0.103 (0.029)** | **4.26e-04** | **-0.127 (0.034)** | **1.81e-04** |
|  |  | Left Lateral occipital | -0.076 (0.029) | 9.51e-03 | **-0.108 (0.034)** | **1.53e-03** |
|  |  | Left Lingual | -0.004 (0.029) | 8.96e-01 | **-0.099 (0.034)** | **3.61e-03** |
|  |  | Left Medial orbitofrontal | 0.057 (0.029) | 5.39e-02 | **0.107 (0.034)** | **1.68e-03** |
|  |  | Left Posterior cingulate | **-0.124 (0.029)** | **2.28e-05** | **-0.166 (0.034)** | **9.57e-07** |
|  |  | Left Supramarginal | -0.01 (0.029) | 7.27e-01 | **-0.097 (0.034)** | **4.50e-03** |
|  |  | Left Insula | **-0.083 (0.029)** | **4.67e-03** | 0.015 (0.034) | 6.62e-01 |
|  |  | Right Caudal anterior cingulate | **-0.09 (0.029)** | **2.02e-03** | -0.035 (0.034) | 3.09e-01 |
|  |  | Right Caudal middle frontal | -0.044 (0.029) | 1.37e-01 | **-0.107 (0.034)** | **1.71e-03** |
|  |  | Right Entorhinal | **-0.114 (0.029)** | **9.69e-05** | -0.054 (0.034) | 1.14e-01 |
|  |  | Right Isthmus cingulate | -0.058 (0.029) | 4.88e-02 | **-0.091 (0.034)** | **7.40e-03** |
|  |  | Right Medial orbitofrontal | **0.087 (0.029)** | **2.95e-03** | **0.104 (0.034)** | **2.19e-03** |
|  |  | Right Middle temporal | **-0.1 (0.029)** | **6.34e-04** | -0.023 (0.034) | 4.92e-01 |
|  |  | Right Posterior cingulate | -0.049 (0.029) | 9.75e-02 | **-0.103 (0.034)** | **2.39e-03** |
| Neglect | Male | Right Middle temporal | -0.146 (0.036) | 6.82e-05 | 0.018 (0.041) | 6.65e-01 |
|  | Female | Left Lingual | -0.027 (0.029) | 3.64e-01 | **-0.124 (0.034)** | **2.49e-04** |
|  |  | Left Posterior cingulate | -0.057 (0.029) | 5.07e-02 | **-0.111 (0.034)** | **1.04e-03** |
|  |  | Right Entorhinal | **-0.1 (0.029)** | **6.55e-04** | -0.002 (0.034) | 9.54e-01 |

*Childhood Abuse.* In young adult males, greater childhood abuse severity was significantly associated with *thinner* cortical regions in the bilateral isthmus cingulate, right banks of the superior temporal sulcus, right middle temporal, left lateral occipital, and right supramarginal areas and, conversely, *thicker* bilateral medial and lateral orbitofrontal cortices. No associations between CM and CT deviations were found in older adult males (Fig. 2A; Table 2). In young adult females, greater childhood abuse severity was associated with *thinner* bilateral caudal anterior cingulate, left inferior parietal, left isthmus cingulate, left posterior cingulate, left insula, right entorhinal, and right middle temporal regions, but *thicker* right medial orbitofrontal cortex. Older adult females showed partially similar results, with greater childhood abuse severity associated with *thinner* bilateral isthmus, posterior, and caudal anterior cingulate regions, left lateral occipital, lingual, and supramarginal areas, but *thicker* bilateral medial orbitofrontal cortices and left inferior temporal cortex (Fig. 2A; Table 2).

*Childhood Neglect.* The effects of childhood neglect on CT deviation scores were not as pronounced or as widespread compared to those observed with abuse. In young adult males, more severe childhood neglect was associated with *thinner* right middle temporal cortex. No significant associations were found in older adult males. In young adult females, more severe childhood neglect was associated with *thinner* right entorhinal cortex. In older adult females, more severe childhood neglect was associated with *thinner* left posterior cingulate and lingual gyrus (Fig. 2B; Table 2).

### Associations Between Surface Area Deviation Scores and ELA

Our analyses revealed significant associations between CM and deviations from normative surface area (SA) in adults, with patterns varying by age, sex, and CM type (Fig. 3; Table 3). Consistent with our previous analyses, no significant associations were found between CM and SA deviation scores in the pediatric cohort.

**Figure 3.**
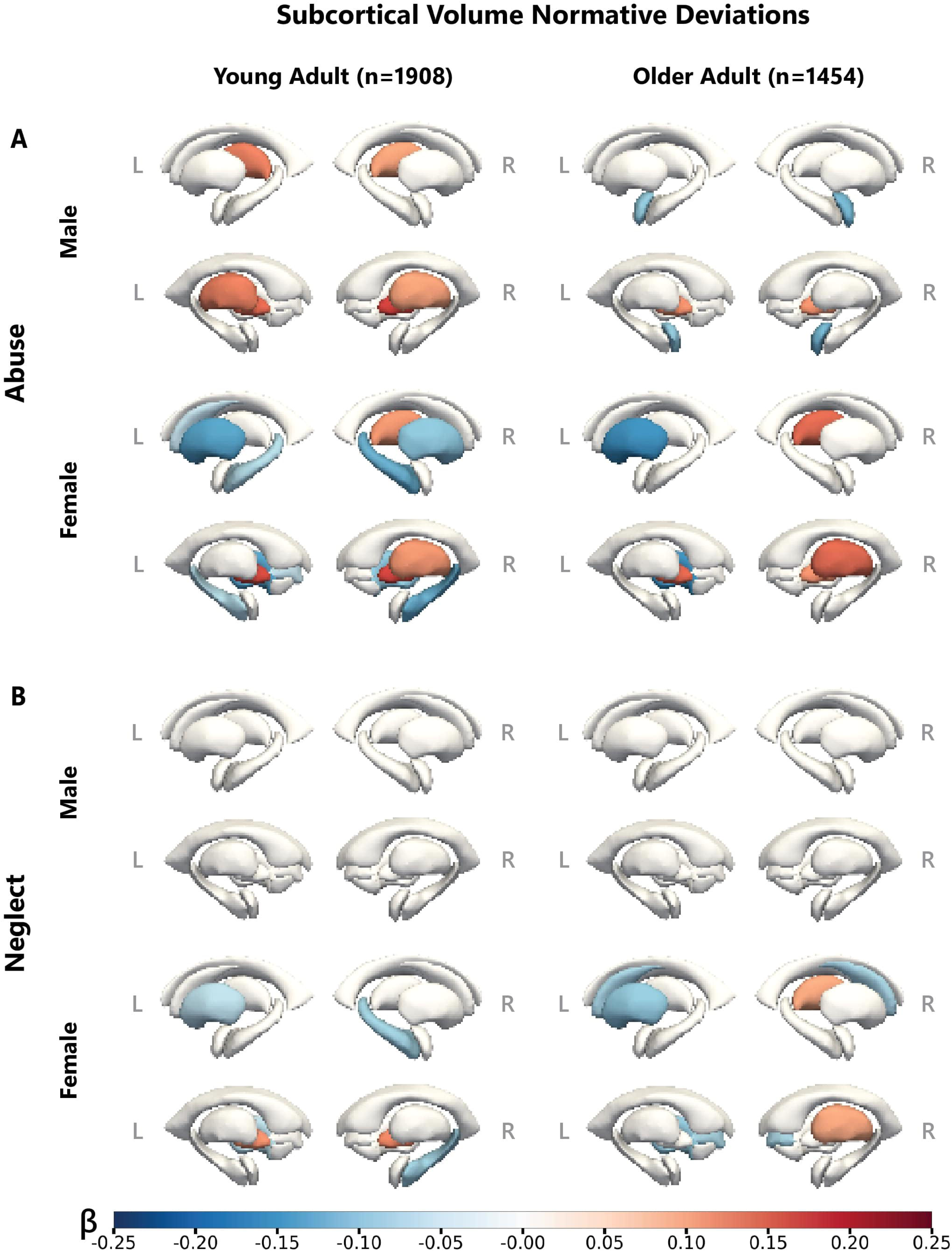
Deviations from Normative Surface Area Associated with Childhood Trauma. **A.** Normative deviations in surface area associated with abuse in males and females; **B.** Normative deviations in surface area associated with neglect in males and females. β represents the standardized effect size from the GLM (FDR corrected *q*<.05). Results from the pediatric cohort are not displayed as there were no significant associations between CTQ and subcortical volumes in this age cohort.

**Table 3.**
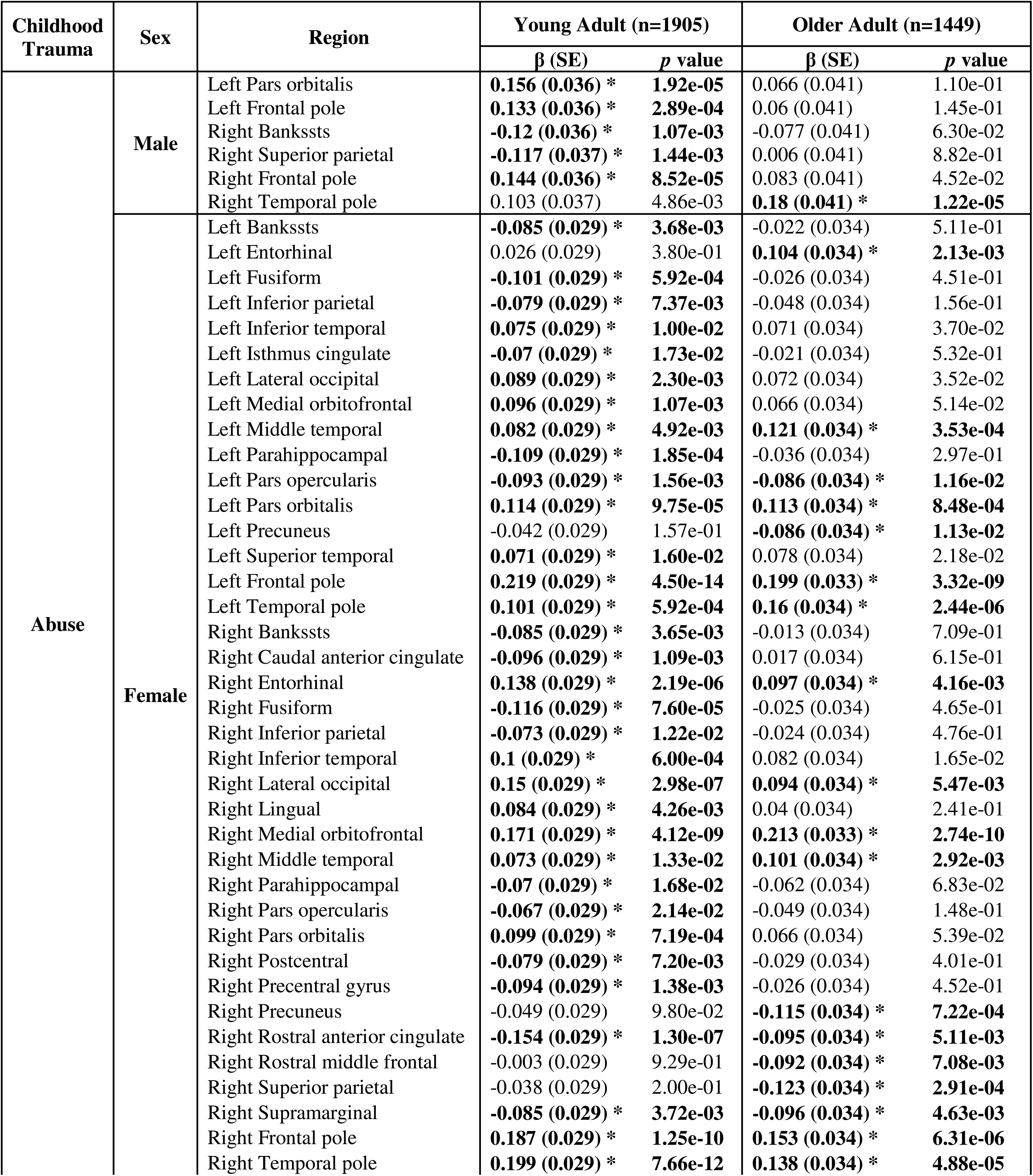
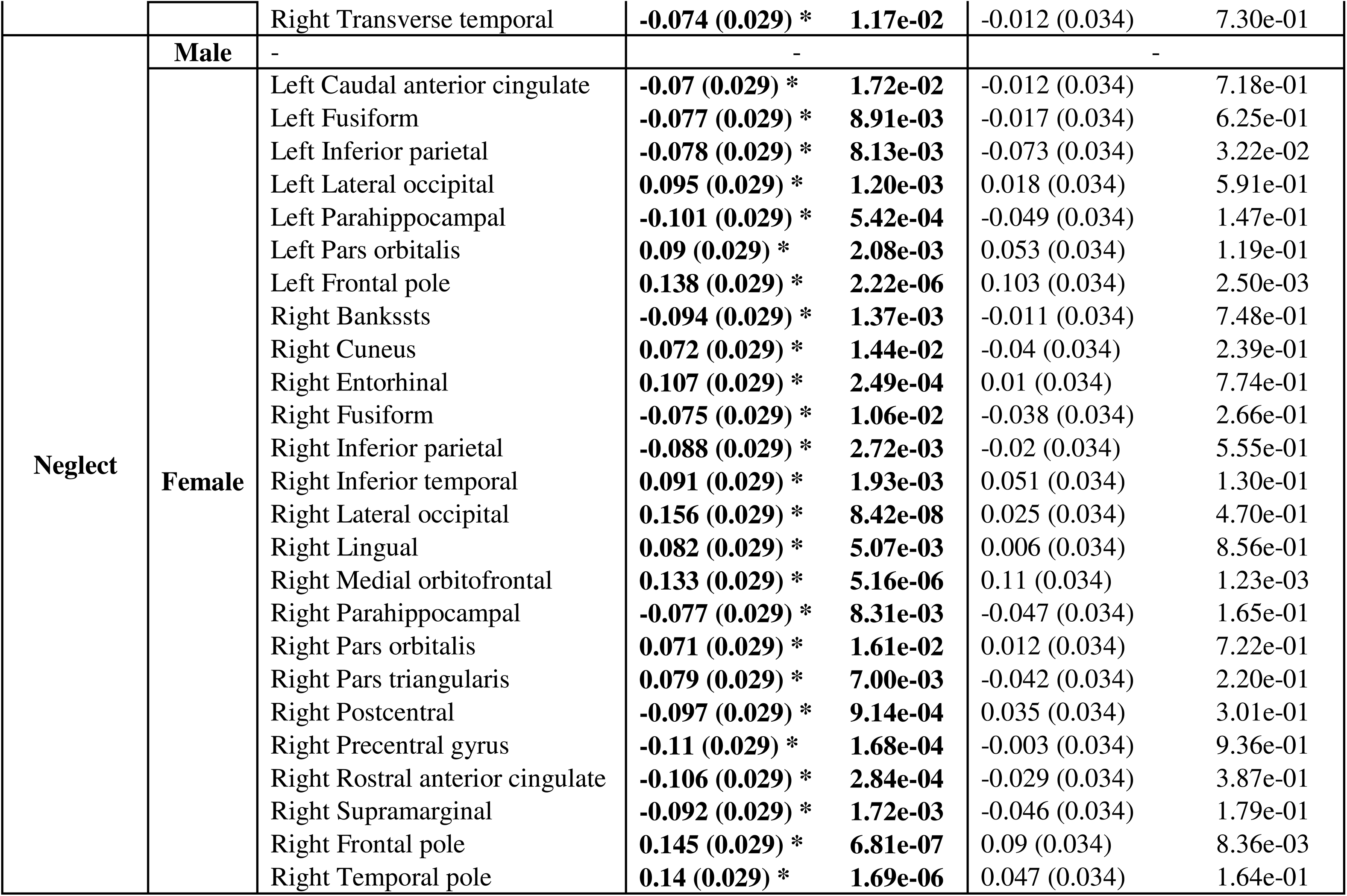
Deviations from Normative Surface Area Associated with Childhood Trauma. P-values are **t**hresholded to show regions with normative values significantly associated with childhood trauma in at least one age cohort. See the full results table of all regions and cohorts in Supplementary Table S4. Significant regions (*q*<.05) are indicated by bolded effect sizes and *p* values. β represents the standardized effect size from the GLM. Results from the pediatric cohort are not displayed as there were no significant associations between CTQ and subcortical volumes in this age cohort.

*Childhood Abuse.* In young adult males, more severe childhood abuse was associated with *smaller* SA in the right banks of the superior temporal sulcus and right superior parietal region but *larger* SA in the bilateral frontal pole. In older adult males, more severe childhood abuse was associated with *smaller* SA of the temporal pole (Fig. 3A; Table 3). In young adult females, more severe childhood abuse was associated with *larger* SA in several regions, including the banks of the left superior temporal sulcus, left isthmus of the cingulate, and bilateral fusiform, inferior parietal, and parahippocampal areas, but *larger* SA in bilateral occipital regions, medial orbitofrontal cortex, pars orbitalis, and the frontal and temporal poles. This pattern persisted in older adult females, although to a lesser extent (Fig. 3A; Table 3).

*Childhood Neglect.* The SA deviations associated with childhood neglect were not as pronounced or as widespread as the effects that we observed with abuse. In males, no significant associations between childhood neglect and SA deviation were detected. In females, associations were only found in young adults: greater severity of childhood neglect was associated with *smaller* SA in bilateral cingulate cortex, parietal lobes, and fusiform regions, as well as *larger* SA in lateral occipital regions, orbitofrontal cortices, and the temporal pole (Fig. 3B; Table 3).

### Sex and Age Differences in Brain-CM Associations

Fisher’s exact tests revealed young adults demonstrated stronger associations than older adults across all three morphometric phenotypes and both types of CM exposure. Additionally, females generally showed stronger associations with CM than males, particularly for SA measures (Table S5).

### Sensitivity Analyses

*Diagnostic Effects.* To examine how clinical status affects associations between CM and brain deviation scores, we analyzed: (1) CM effects adjusting for diagnostic status, and (2) diagnostic effects adjusting for CM (Supplementary Methods S5; Table S6). We statistically adjusted for diagnosis as a dichotomous variable (patient vs. control), grouping MDD and PTSD patients together. After adjusting for diagnosis, most ELA-brain associations remained consistent with our primary findings across age cohorts (Table S6). Childhood abuse associations persist across young and older adult females, while certain childhood neglect associations unique to older adult females were attenuated after accounting for diagnosis.

Diagnosis itself was associated with widespread cortical thickness deviations in pediatric and older adult female cohorts, particularly in bilateral frontal, temporal, and occipital regions (Table S6). For SA, diagnostic adjustment eliminated significant associations in young adults but not in the older adult cohort. Overall, most ELA-brain associations persist independent of psychiatric diagnosis, though MDD/PTSD diagnoses may contribute to additional brain morphometric deviations unrelated to CM exposure.

*Site-Specific Effects.* To assess site-specific influences on our findings, we performed leave-one-out analysis across all sites. Results remained highly consistent across the three morphometric phenotypes, childhood trauma types, sexes, and age cohorts (Table S7). The pediatric cohort showed greater variability in *p*-values due to smaller sample sizes, yet consistently yielded non-significant results. Young adult and older adult cohorts demonstrated robust associations between childhood trauma and brain morphometry across sites, with key subcortical and cortical regions maintaining significance despite site variability.

*Harmonization Effect.* ComBat-GAM harmonization yielded patterns that were similar to our main results (Fig. S4). Key findings remained consistent: in young adult females, both abuse and neglect were associated with smaller right hippocampal volumes (β=-.09, *q*<.05); in older adult males, abuse was associated with thicker left superior parietal cortex (β=.14, *q*<.05); and in young adult males, neglect was associated with thinner right middle temporal regions (β=-.14, *q*<.05). These consistent findings, despite relatively small effect sizes, support the robustness of our results.

*Power Analysis for Pediatric Cohort.* Bootstrap down-sampling revealed only 3.4% of significant associations in adult cohorts would be detectable with the pediatric sample size, well below the typical threshold for power (80%). Only the effects of childhood abuse on surface area in specific frontal regions were sufficiently powered. The result from this analysis indicates that while null findings in children likely reflect limited statistical power rather than definitive age-related differences, the effects we identified in certain frontal regions are nonetheless robust enough to be detected with smaller samples (Supplementary Results S1).

## DISCUSSION

In this largest investigation to date of the effects of CM on brain morphometry, we found widespread neuroanatomical alterations that varied by age, sex, severity of ELA, and type of ELA. The most pronounced effects were found for childhood abuse, which showed extensive effects across several subcortical and cortical regions, particularly among young adult females (although many of these patterns persisted to a lesser degree in older adult females). Young adult females also exhibited the most pronounced effects of neglect, although these effects were limited to deviations in surface area and appeared to be unique to cingulate regions (among young adult females, both abuse and neglect were associated with smaller right entorhinal cortex, smaller surface area in fusiform gyrus and inferior parietal regions, larger surface area of lateral occipital and medial orbitofrontal cortices, as well as larger temporal poles). Effects of neglect on male brain morphometry were observed only in young adult males and were associated only with smaller right middle temporal cortex (a pattern also found in young adult males and females exposed to greater childhood abuse). Both young adult females and males exposed to abuse exhibited larger volumes in the bilateral thalamus and pallidum, as well as smaller cortices in isthmus cingulate and larger medial orbitofrontal cortices.

Broadly, these distinct patterns support longstanding theories and emerging evidence that different forms of CM differentially shape neurocircuits implicated in affective, cognitive, and socioemotional processing [16, 35] and that many of these effects are sex-specific [29, 55] and transdiagnostic [56–58]. Our findings revealed bidirectional relationships between CM and brain morphometry, with some regions showing larger structural deviations and others showing smaller (i.e., more negative) deviations within the same age cohort. These opposing patterns likely reflect complex reorganizational changes across neural circuits rather than uniform effects, potentially resulting from compensatory adaptation and learning during development [59–61]. Although we did not detect significant associations in the pediatric cohort, this null finding is likely attributable to limited statistical power, based on our power analysis.

Interestingly, CM effects on brain morphometry in young adults were primarily observed in association cortices [62]. These higher-order regions, which typically undergo more protracted development, showed notable negative z-score deviations in response to early adversity. This pattern aligns with the hierarchical nature of cortical development, where association cortices exhibit more protracted development and extended plasticity compared to primary sensory regions [62–64]. This extended period of plasticity may make association cortices more vulnerable to the effects of ELA, especially during adolescence and young adulthood when these regions are still undergoing significant refinement [65, 66]. Our finding of more widespread associations between CM and surface area in young adults, with both measures affected in older adults, reflects their distinct developmental properties— cortical surface area is more heritable and established earlier, while thickness remains more responsive to environmental influences throughout life [67, 68]. Furthermore, the observed alterations in association cortices—critical for higher-order functions including emotion regulation, executive control, and social cognition [56, 69]—may help explain the substantially increased risk for stress-related psychopathology associated with ELA, including major depressive disorder (OR=2.7) [70] and posttraumatic stress disorder (OR=4.4) [71, 72]. Nonetheless, our cross-sectional design limits our ability to make strong developmental claims, highlighting the need for longitudinal studies to establish the temporal sequence of the effects we observed in our analyses [73, 74].

The more pronounced effects in females, particularly during young adulthood, suggest that there are important sex-specific neural vulnerabilities to CM to consider within a *sensitive period* framework.

While additional data are needed to corroborate this formulation, the notable female vulnerability may reflect interactions between CM and sex-specific neurodevelopmental trajectories, potentially mediated by gonadal hormones and their organizational effects on brain development [16, 75, 76]. Young women between the ages of 18 and 34 experience relatively stable hormonal profiles [77]. This period of hormonal stability may create conditions where the effects of prior CM can consolidate and leave more enduring neurobiological impacts [78]. Additionally, sociocultural factors, including differential exposure to subsequent stressors and varying societal expectations and pressures, may contribute to sex differences in the manifestation of adversity-related brain alterations [32, 76, 79]. It may also be helpful to consider other influences, such as the potential "normalizing" effect of treatment interventions in older individuals, which could diminish observable neuroanatomical differences over time [34].

While we found no significant associations between CM and structural brain deviation scores in our pediatric cohorts, our power analysis indicates this is primarily due to limited statistical power. Our bootstrap down-sampling analysis revealed that most significant associations observed in young adults would not be detectable in our pediatric sample due to sample size (*n*=349). Nevertheless, the associations we observed between CM and structural brain deviation scores in adults demonstrate the existence of normative deviations in brain structure related to childhood adversity during adult stages, which aligns with a sensitive period model of ELA [19, 21]. However, given the power constraints in our pediatric sample, we cannot make definitive conclusions regarding the incubation model that would predict different effect magnitudes across developmental stages. Similarly, we cannot adequately evaluate accumulation models of ELA, which propose that the impact of early adversity on brain structure progressively increases throughout development. Future research, ideally using longitudinal study designs, will need to ensure adequate and consistent statistical power across age cohorts to properly characterize whether and, if so, how there are differential impacts of CM on brain morphometry throughout the lifespan.

Our study has several limitations that warrant additional research studies to address. First, our data were cross-sectional, so we could not characterize causal effects of CM exposure on the brain as a function of age, as we were not directly assessing within-person neurodevelopmental trajectories. Second, the Childhood Trauma Questionnaire, while widely adopted and well validated, does not capture all forms of ELA, such as housing instability, toxicant exposure, and neighborhood violence, potentially omitting significant contributors to brain development [80, 81]. Third, our power analysis demonstrated that the relatively smaller pediatric sample size was insufficient to detect the subtle effects observed in adults, limiting our ability to draw definitive conclusions about developmental timing; and we could not fully account for potential confounding factors such as socioeconomic status, education level, symptom severity, and medication use, which could independently influence brain structure but were confounded with diagnosis and study site in our dataset. Additionally, the heterogeneity in our sample, including variations in the timing, chronicity, and combination of CM types, may obscure more nuanced associations between specific adversities and neurobiological outcomes. Future research should employ longitudinal designs to elucidate developmental trajectories and should incorporate comprehensive assessments of ELA, including a broader range of adversities and detailed environmental contexts. Indeed, our findings underscore the importance of incorporating CM measures in developmental research across psychiatric disorders—including anxiety, psychosis, and personality disorders—where early adversity has been implicated in etiology [74, 82, 83] but its neurobiological impact remains inadequately characterized. Furthermore, consideration of additional variables such as genetic and epigenetic factors [15], resilience mechanisms [75], and interactions between environmental influences [84] will be crucial to advance our understanding of the complex pathways linking CM to brain development.

Despite these limitations, our findings have important clinical implications for research and treatment approaches. Individuals with MDD or PTSD who experienced CM show distinct patterns of brain normative deviation compared to those without CM exposure, suggesting potentially different neurobiological pathways to these disorders. This neurobiological heterogeneity may partially explain the mixed response to standard treatments often observed in clinical practice. While our results do not directly address treatment resistance, they suggest the value of assessing CM history when developing treatment plans, as recognizing these distinct neurobiological patterns may help inform more comprehensive approaches to care for trauma-exposed individuals. Future research is needed to determine whether these distinct brain morphometric patterns predict differential responses to pharmacological and psychological interventions.

## CONCLUSION

Our investigation reveals that childhood maltreatment has widespread effects on brain structure, with particularly pronounced effects on association cortices in young adult females. The distinct patterns observed for abuse versus neglect, along with the sex-dependent effects, underscore the complexity of how early adversity shapes brain development, especially during periods of heightened susceptibility. These findings highlight the potential importance of early intervention, even in the potential absence of immediate structural brain alterations. Adolescence and young adulthood may represent vital points for intervention, particularly for females with experiences of childhood maltreatment.

### Data Sharing Statement

The datasets analyzed during the current study are not publicly available due to site restrictions but data may be available from the corresponding sites on reasonable request.

### Conflicts of Interest Disclosures

CA served as a consultant and on advisory boards for Douglas Pharmaceutical, Freedom Biosciences, Aptinyx, Genentech, Janssen, Psilocybin Labs, Lundbeck, Guidepoint, and FSV7. He also received royalties from a patent for using mTORC1 inhibitors to augment the effects of antidepressants.

IHR reports research funding from BI Pharma on precision psychiatry. RH is a consultant for Jazz Pharmaceuticals.

LL reports unpaid membership on the Scientific Committee for the International Society for the Study of Trauma and Dissociation (ISSTD) and spousal IP payments from Vanderbilt University for technology licensed to Acadia Pharmaceuticals and spousal private equity in Violet Therapeutics unrelated to the present work.

EAO reports employment at Crisis Text Line.

KR has performed scientific consultation for Bioxcel, Bionomics, Acer, and Jazz Pharma; serves on Scientific Advisory Boards for Sage, Boehringer Ingelheim, Senseye, and the Brain Research Foundation, and he has received sponsored research support from Alto Neuroscience.

AE reports salary and equity from Alto Neuroscience, equity from Mindstrong Health and Akili Interactive.

TY reports a clinical trials agreement that provides funds to UCSF by NeuroQore, and a portion of these funds to UCSF have supported TTY’s effort on paper(s) with NeuroQore. None of the funds from NeuroQore supported Dr Yang’s effort on this current paper.

All other authors do not have any conflicts of interest to declare.

## Acknowledgements

ZQL is supported by Fonds de Recherche du Québec – Nature et Technologies (FRQNT); AH is supported by R01 MH111671 and VISN6 MIRECC; CA is supported by the Beth K and Stuart Yudofsky Chair in the Neuropsychiatry of Military Post Traumatic Stress Syndrome; NA is supported by the Deutsche Forschungsgemeinschaft (DFG, grant to SFB/TRR 393 – Project-ID 521379614).

JUB is supported by NIMH MH106998; UD is supported by the German Research Foundation (DFG) grant FOR2107 DA1151/5-1, DA1151/5-2, DA1151/9-1, DA1151/10-1, DA1151/11-1 to UD; SFB/TRR 393, project grant no 521379614) and the Interdisciplinary Center for Clinical Research (IZKF) of the medical faculty of Münster; TdC is supported by R01 MH106574; DD is supported by the National Institute for Health and Care Research (NIHR) Maudsley Biomedical Research Centre, South London and Maudsley NHS Trust; NF is supported by MH101380 and MH111671-01A1; CHF is supported by the National Institute for Health and Care Research (NIHR) Maudsley Biomedical Research Centre (Grant JR 4), and the International Psychoanalytical Association (IPA 158102845); EMG is supported by R01 NS140256; IG is supported by NIMH R37 101495; RGM is supported by the German Federal Ministry of Education and Research (BMBF: 01 ZX 1507, “PreNeSt – e:Med”); NAG is supported by the Bill & Melinda Gates Foundation [OPP 1017641]; IHR is supported by MH102634 and the U.S. National Center for PTSD; RH is supported by NIMH R01 MH132221 and R01 MH128371; SLB is supported by R01 MH126814 and U01 MH121737; AB is supported by JDRF (3-SRA-2019-836-M-B) and the Department of Defense (W81XWH-18-1-0230); LG is supported by R01 MH117229, R01 MH118181, R61 AT009864-01A1, R33 AT009864, and the Beckman Foundation; JCS is supported by partial grants from NIMH (R01 MH12345), SUNOVION (Research Grant), and Mind Medicine (Research Grant); DJS is supported by the South African Medical Research Council (SAMRC); FS is supported by the German Research Foundation (DFG, SFB/TRR 393, project grant no. 521379614); BSJ is supported by NIMH R01 MH131532; JT is supported by the Canadian Institutes of Health Research (CIHR) and the Canadian Institute for Military and Veteran Health Research (CIMVHR); SIT is supported by R01 MH121806, R01 MH129742, and R01 AG058854 (to PMT); LLvdH is supported by the South African Medical Research Council (SAMRC) MRC-RFA-IFSP-01-2013/SHARED ROOTS; XW is supported by R21 MH098198-01 and the ProMedica Translational Research Stimulation Award; HW is supported by Wellcome Trust funding (220857/Z/20/Z); TY is supported by the National Center for Complementary and Integrative Health (1R61 AT009864-01A1 and R33 AT009864); XZ is supported by K01 MH122774; PMT is supported by R01 MH121806, R01 MH129742, and R01 AG058854; LH is supported by a Veni Award (grant number 09150162210201) from the Dutch Research Council (NWO); LS is supported by NIH R01 MH129742 and by a Dame Kate Campbell Fellowship at the University of Melbourne; TCH is supported by K01 MH117442 (TIGER), the Klingenstein Third Generation Foundation (Child and Adolescent Depression Fellow Award), the Stanford Maternal Child and Health Institute (Early Career Award and K Support Award), the Stanford Center for Cognitive and Neurobiological Imaging (Seed Grant), the Ray and Dagmar Dolby Family Fund, and NIH (R01 MH127176, R21 MH130817).

We thank Erin O’Leary, Geoffrey May, Orren Ravid, and Julia Herzog for their contributions to data collection.

## ENIGMA MDD and PTSD Working Groups Authors

Haley R. Wang^¹,²^, Zhen-Qi Liu^³^, Elena Pozzi, Ahmed Hussain, Priyanka Sigar, Chadi Abdallah, Nina Alexander^¹^, Justin Baker^¹¹,¹²^, Jochen Bauer^¹,³^, Jennifer Urbano Blackford^¹,¹^, Thomas Boeck^¹^, Josh Cisler^¹^, Colm G. Connolly^¹^, Andrew Cotton^¹^, Judith Daniels^¹,²^, Udo Dannlowski^²¹^, Maria Densmore^²²,²,³^, Terri deRoon-Cassini^²,²^, Danai Dima^²,²^, Stefan Du Plessis^²,²^, Amit Etkin^³^, Negar Fani^³,¹^, Lukas Fisch^²¹^, Jacklynn Fitzgerald^³,²^, Thomas Frodl^³,³^, Cynthia H.Y. Fu^³,,³^, Ali Saffet Gonul^³^, Evan M. Gordon^³^, Ian Gotlib^³^, Roberto Goya-Maldonado^³^, Nynke A. Groenewold, Dominik Grotegerd^²¹^, Tim Hahn^²¹^, J. Paul Hamilton^¹^, Ilan Harpaz-Rotem^²^, Courtney C. Haswell^³^, Ryan Herringa, Jonathan Ipser, Neda Jahanshad, Tanja Jovanovic, Milissa Kaufman^¹¹,¹²^, Tilo Kircher^¹^, Maximilian Konowski^²¹^, Sheri-Michelle Koopowitz, Axel Krug, John Krystal^³,^, Ruth Lanius, Christine Larson, Amit Lazarov^¹,²^, Lauren Lebois^¹¹,¹²^, Elisabeth Leehr^²¹^, Ifat Levy^³^, Meng Li, Antje Manthey, Antje Manthey, Adi Maron-Katz, Andrew McIntosh, Katie McLaughlin, Susanne Meinert^²¹,¹^, Rajendra Morey^²^, Benson Mwangi^³,^, Steven M. Nelson, Igor Nenadić^¹^, Yuval Neria^²^, Richard Neufeld, Amar Ojha, Bunmi Olatunji^¹^, Elizabeth A. Olson^²^,Nils Opel^²¹,³^, Matthew Peverill, Kerry Ressler, Marisa Ross, Isabelle Rosso, Matthew Sacchet, Kelly Sambrook, Christian Schmahl^¹^, Soraya Seedat^²,³^, Martha E. Shenton, Maurizio Sicorello^¹^, Anika Sierk, Jair C. Soares^³,^, Dan J. Stein, Frederike Stein^¹^, Murray B. Stein, Benjamin Suarez-Jimenez, Jean Théberge^²²^, Sophia I. Thomopoulos, Carissa Tomas^²^, Paula Usemann^¹^, Leigh L. van den Heuvel^²,³^, Sanne van Rooij^³,¹^, Martin Walter, Henrik Walter^¹^, Xin Wang^²^, Heather Whalley, Nils R. Winter^²¹^, Mon-Ju Wu^³,^, Hong Xie^²^, Tony Yang^³^, Xi Zhu, Sigal Zilcha-Mano, Giovana B. Zunta- Soares^³,^, Sophia Frangou, Paul M. Thompson, Laura Han^¹^, Lauren Salminen, Delin Sun^¹¹,¹²,¹³^, Jennifer S. Stevens^¹,¹^, Lianne Schmaal, Tiffany C. Ho^¹^

## Affiliations of ENIGMA MDD and PTSD Working Groups Authors

1. Department of Psychology, University of California, Los Angeles, Los Angeles, CA, USA
2. Semel Institute for Neuroscience and Human Behavior, University of California, Los Angeles, Los Angeles, CA, USA
3. Montréal Neurological Institute, McGill University, Montréal, QC, Canada
4. Centre for Youth Mental Health, The University of Melbourne, Melbourne, Australia
5. Orygen, The National Centre of Excellence in Youth Mental Health, Parkville, Australia
6. Brain Imaging and Analysis Center, Duke University, Durham, NC, USA
7. Department of Psychiatry and Behavioral Sciences, University of California, Los Angeles, Los Angeles, CA, USA
8. Department of Psychiatry, Baylor College of Medicine, Houston, TX, USA
9. Department of Psychiatry, Yale University, New Haven, CT, USA
10. Department of Psychiatry, University of Marburg, Marburg, Germany
11. McLean Hospital, Belmont, MA, USA
12. Harvard Medical School, Boston, MA, USA
13. Clinic for Radiology, University of Münster, Münster, Germany
14. Munroe-Meyer Institute, University of Nebraska Medical Center, Omaha, NE, USA
15. Department of Psychiatry and Behavioral Sciences, Vanderbilt University Medical Center, Nashville, TN, USA
16. University of Texas at Austin, Austin, TX, USA
17. Department of Biomedical Sciences, Florida State University, Tallahassee, FL, USA
18. Department of Neurology, Division of Cognitive Neuroscience, Johns Hopkins University School of Medicine, Baltimore, MD, USA
19. Department of Clinical Psychology, University of Groningen, The Netherlands
20. TraumaCentrum Beilen, GGZ Drenthe, Beilen, The Netherlands
21. Institute for Translational Psychiatry, University of Münster, Münster, Germany
22. Department of Psychiatry, Western University, London, ON, Canada
23. Imaging Division, Lawson Research Institute, London, ON, Canada
24. Division of Trauma and Acute Care Surgery, Department of Surgery, Medical College of Wisconsin, WI, USA
25. Comprehensive Injury Center, Medical College of Wisconsin, WI, USA
26. Department of Psychology, City St George’s, University of London, London, UK
27. Department of Neuroimaging, Institute of Psychiatry, Psychology and Neuroscience, King’s College London, London, UK
28. Department Psychiatry, FMHS, Stellenbosch University, Tygerberg, Cape Town, South Africa
29. Genomics of Brain Disorders Research Unit, South African Medical ResearchCouncil/Stellenbosch University, Cape Town, South Africa
30. Stanford University, Stanford, CA, USA
31. Department of Psychiatry and Behavioral Sciences, Emory University, Atlanta, GA, USA
32. Department of Psychology, Marquette University, Milwaukee, WI, USA
33. Department of Psychiatry, Psychotherapy and Psychosomatics, University Hospital Aachen, Aachen, Germany
34. Centre for Affective Disorders, Institute of Psychiatry, Psychology and Neuroscience, King’s College London, London, UK
35. Department of Psychology, University of East London, London, UK
36. Department of Psychiatry, Ege University School of Medicine, Izmir, Turkey
37. Department of Radiology, Washington University School of Medicine, St. Louis, MO, USA
38. Department of Psychology, Stanford University, Stanford, CA, USA
39. Department of Psychiatry and Psychotherapy, University Medical Center Göttingen, Göttingen, Germany
40. Department of Psychiatry & Neuroscience Institute, University of Cape Town, Cape Town, South Africa
41. Department of Biological and Medical Psychology, University of Bergen, Bergen, Norway
42. Departments of Psychiatry and of Psychology, Yale University, New Haven, CT, USA
43. Clinical Neuroscience Division, National Center for PTSD, VA Connecticut Healthcare System, West Haven, CT, USA
44. Department of Psychiatry, University of Wisconsin School of Medicine & Public Health, Madison, WI, USA
45. Mark and Mary Stevens Neuroimaging and Informatics Institute, Keck School of Medicine of USC, University of Southern California, Marina del Rey, CA, USA
46. Department of Psychiatry and Behavioral Neurosciences, Wayne State University, Detroit, MI, USA
47. Department of Psychiatry and Psychotherapy, University Hospital Bonn, Bonn, Germany
48. Department of Psychiatry, Yale University School of Medicine, New Haven, CT, USA
49. Department of Neuroscience, Western University, London, ON, Canada
50. Department of Psychology, University of Wisconsin-Milwaukee, Milwaukee, WI, USA
51. School of Psychological Sciences, Tel Aviv Univeristy, Tel Aviv, Israel
52. Irving Medical Center, Columbia University, New York, NY, USA
53. Departments of Comparative Medicine, Neuroscience, and Psychology, Yale University, New Haven, CT, USA
54. Department of Psychiatry and Psychotherapy, University Hospital Jena, Jena, Germany
55. Health and Medical University Potsdam, Potsdam, Germany
56. Charité Universitätsmedizin Berlin Campus Charite Mitte: Charité Universitätsmedizin Berlin, Berlin, Germany
57. Department of Psychiatry and Behavioral Sciences, Stanford University, Stanford, CA, USA
58. Centre for Clinical Brain Sciences, University of Edinburgh, Edinburgh, Scotland, UK
59. Department of Psychology, Harvard University, Cambridge, MA, USA
60. Ballmer Institute for Children’s Behavioral Health, University of Oregon, Eugene, OR, USA
61. Institute for Translational Neuroscience, University of Münster, Münster, Germany
62. Duke University, Durham, NC, USA
63. Center of Excellence on Mood Disorders, Louis A. Faillace, MD, Department of Psychiatry and Behavioral Sciences, The University of Texas Health Science Center at Houston, Houston, TX, USA
64. School of Behavioral Health Sciences, The University of Texas Health Science Center at Houston, Houston, TX, USA
65. Department of Pediatrics, University of Minnesota, Minneapolis, MN, USA
66. Masonic Institute for the Developing Brain, University of Minnesota, Minneapolis, MN, USA
67. Departments of Psychology and Psychiatry, Neuroscience Program, Western University, London, ON, Canada
68. Department of Psychology, University of British Columbia, Okanagan, Kelowna, BC, Canada
69. Center for Neuroscience, University of Pittsburgh, Pittsburgh, PA, USA
70. Center for the Neural Basis of Cognition, University of Pittsburgh, Pittsburgh, PA, USA
71. Department of Psychology, Nashville, TN, USA
72. Crisis Text Line, New York, NY, USA
73. German Center for Mental Health (DZPG), Berlin, Germany
74. Department of Psychiatry, University of Wisconsin-Madison, Madison, WI, USA
75. Division of Depression and Anxiety Disorders, McLean Hospital, Belmont, MA, USA
76. Department of Psychiatry, Harvard Medical School, Boston, MA, USA
77. Northwestern University, Evanston, IL, USA
78. Meditation Research Program, Department of Psychiatry, Massachusetts General Hospital, Harvard Medical School, Boston, MA, USA
79. University of Washington, Seattle, WA, USA
80. Department of Psychosomatic Medicine and Psychotherapy, Central Institute of Mental Health, Medical Faculty Mannheim, Heidelberg University, Heidelberg, Germany
81. German Center for Mental Health (DZPG), Partner Site Mannheim-Heidelberg-Ulm, Mannheim, Germany
82. Department of Psychiatry, Faculty of Medicine and Health Sciences, Stellenbosch University, Cape Town, South Africa
83. South African Medical Council Unit on the Genomics of Brain Disorders, Department of Psychiatry, Faculty of Medicine and Health Sciences, Stellenbosch University, Stellenbosch, South Africa
84. Department of Psychiatry, Mass General Brigham (MGB), Brigham and Women’s Hospital and Harvard Medical School, Boston, MA, USA
85. SAMRC Unit on Risk & Resilience in Mental Disorders, Department of Psychiatry & Neuroscience Institute, University of Cape Town, Cape Town, South Africa
86. Department of Psychiatry, University of California, San Diego, San Diego, CA, USA
87. The Del Monte Institute for Neuroscience, Department of Neuroscience, University of Rochester School of Medicine and Dentistry, Rochester, NY, USA
88. Imaging Genetics Center, Mark and Mary Stevens Neuroimaging and Informatics Institute, Keck School of Medicine, University of Southern California, Marina del Rey, CA, USA
89. Division of Epidemiology and Social Sciences, Institute of Health and Humanity, Medical College of Wisconsin, WI, USA
90. German Center for Mental Health (DZPG), Mannheim, Germany
91. Division of Mind and Brain Research, Department for Psychiatry, Charité– Universitätsmedizin Berlin, Berlin, Germany
92. Department of Neuroscience and Psychiatry, University of Toledo, Toledo, OH, USA
93. Department of Psychiatry, Division of Child and Adolescent Psychiatry, School of Medicine, University of California, San Francisco, San Francisco, CA, USA
94. Department of Bioengineering, The University of Texas at Arlington, Arlington, TX, USA
95. School of Psychological Sciences, University of Haifa, Haifa, Israel
96. Djavad Mowafaghian Centre for Brain Health, University of British Columbia, Vancouver, BC, Canada
97. Department of Psychiatry, Icahn School of Medicine at Mount Sinai, New York, NY, USA
98. Department of Psychiatry, Amsterdam UMC location Vrije Universiteit Amsterdam, Amsterdam, The Netherlands
99. Amsterdam Neuroscience, Mood, Anxiety, Psychosis, Sleep & Stress program, Amsterdam, The Netherlands
100. Amsterdam Public Health, Mental Health program, Amsterdam, The Netherlands
101. Department of Psychiatry and Behavioral Sciences, School of Medicine, Duke University, Durham, NC, USA
102. Brain Imaging Analysis Center, Duke University, Durham, NC, USA
103. Department of Veteran Affairs Mid-Atlantic Mental Illness Research, Education and Clinical Center, Durham, NC, USA
104. Department of Psychiatry and Behavioral Sciences, Emory University School of Medicine, Atlanta, GA, USA
105. Atlanta VA Medical Center, Atlanta, GA, USA

