## Supplemental material for "Childhood Maltreatment and Deviations from Normative Brain Structure: Results from 3,711 Individuals from the ENIGMA MDD and ENIGMA PTSD"

***Supplemental Information***

**Supplemental Methods**

**Methods S1.** Detailed information on participant recruitment, data collection procedures, and MRI processing for each cohort

**Methods S2.** ENIGMA Consortium's standardized quality control and processing pipelines for MRI data

**Methods S3.** CentileBrain normative modeling framework

**Methods S4.** Sex and age differences in associations between FreeSurfer metrics and CTQ

**Methods S5.** Post-hoc analysis testing effect of diagnosis

**Methods S6.** Testing site-specific effects using leave-one-out cross-validation

**Methods S7.** Results with ComBat-GAM for site harmonization

**Methods S8.** Power analysis for pediatric cohort

**Supplemental Results**

**Figure S1**. Geographical distribution of participating research sites

**Table S1.** ENIGMA sites sample demographics

**Table S2.** ENIGMA sites scan image acquisition and processing

**Table S3.** ENIGMA sites instrument for clinical diagnosis and local study eligibility criteria

**Figure** **S2.** Distribution of age by diagnosis

**Figure** **S3.** Distribution of CTQ scores by age and diagnosis

**Table S4.** Full results of normative deviations associated with CTQ

**Table S5.** Fisher's exact tests on normative deviations between age cohorts and between sex

**Table S6.** Full results of normative deviation-CTQ association controlling for diagnosis

**Table S7.** Leave-one-out analysis results demonstrating the robustness of findings across sites

**Figure S4.** Main results with ComBat-GAM harmonization

**Result S1.** Power analysis for pediatric cohort

**Supplemental References**

This supplementary material has been provided by the authors to give readers additional information about their work.

**Supplemental Methods**

**Methods S1. Detailed information on participant recruitment, data collection procedures, and MRI processing for each cohort**

The current study analyzed data from 25 independent sites spanning 8 countries, assembled through the ENIGMA Major Depressive Disorder (MDD) and Posttraumatic Stress Disorder (PTSD) Working Groups (see Supplementary Results S1 and Supplementary Table S1 for detailed demographics and clinical characteristics of each cohort). Participants were recruited according to site-specific inclusion and exclusion criteria, standardized across sites through structured diagnostic interviews (e.g., SCID, MINI) based on DSM-IV or ICD-10 criteria. All neuroimaging data were acquired and processed following ENIGMA Consortium standardized imaging protocols and rigorous quality assurance procedures, ensuring consistency and comparability across datasets (see Supplementary Methods S2, Supplementary Table S2, and Figure S1 for scanner details and imaging protocols).

Notably, among all sites contributing to the two ENIGMA work groups and consenting to inclusion in this study, we only included sites with complete structural MRI and Childhood Trauma Questionnaire–Short Form (CTQ-SF) data. Sites lacking data from the CTQ-SF or that used a different version of the CTQ were not included. Additionally, some sites provided only the CTQ total score without individual item scores or subscale scores for abuse and neglect, and thus could not be included in our intended analysis design. The final sample comprised 25 sites from the two work groups. Eight subjects were included in the subcortical analysis but excluded from the cortical analysis due to missing cortical data.

**Methods S2. ENIGMA Consortium's standardized quality control and processing pipelines for MRI data**

The Enhancing NeuroImaging Genetics through Meta-Analysis (ENIGMA) Consortium has established comprehensive, standardized pipelines for processing and quality control (QC) of neuroimaging data. These protocols are designed to ensure reproducibility and comparability across diverse datasets and research sites. ENIGMA's structural MRI protocols utilize FreeSurfer for cortical and subcortical segmentation, followed by rigorous QC procedures. These include visual inspection of segmentation outputs and the use of automated scripts to identify and correct errors. Specialized protocols have been developed for longitudinal analyses, hippocampal subfield segmentation, and cerebellar volumetrics, each accompanied by tailored QC guidelines (1-3). These standardized pipelines and QC procedures developed by the ENIGMA Consortium are publicly available and have been widely adopted in the neuroimaging community, promoting data harmonization and facilitating large-scale collaborative studies.

**Methods S3.** CentileBrain normative modeling framework

CentileBrain is a normative modeling framework specifically developed to quantify deviations in brain morphometric measures relative to normative patterns observed across the lifespan. Ge et al. (2024) (4) systematically benchmarked and optimized this framework using a comprehensive dataset comprising 37,407 healthy individuals aged between 3 and 90 years from diverse geographical regions. Eight statistical modeling algorithms were empirically compared, with multivariate fractional polynomial regression (MFPR) identified as optimal based on a balance of predictive accuracy and computational efficiency. Optimal performance was achieved with non-linear fractional polynomials for age and linear adjustments for global neuroimaging measures (intracranial volume for subcortical volumes, mean cortical thickness for regional cortical thickness, and total cortical surface area for regional cortical surface area).

Notably, CentileBrain constructs separate normative models for females and males, ensuring that the resulting z-scores accurately reflect sex-specific differences in brain morphology. The optimal model parameters—identified through benchmarking of multiple algorithms and covariate optimization—were applied to each regional measure in our sample. The potential overlap between training samples in these models and healthy controls in our dataset strengthens alignment with normative trajectories rather than introducing bias.

The CentileBrain models were applied to the entire pooled sample of patients and controls to generate a z-score for each morphometric measure in each participant. A z-score quantifies the deviation of the observed value from the population mean and is computed by subtracting the estimated value from the raw value of that measure, and then dividing the difference by the root mean square error of the model. All z-scores are specific to the sex and age of each participant. While the CentileBrain algorithms include site harmonization based on Combat-GAM (5), some of the sites in our analyses lacked a cohort of healthy controls, likely resulting in meaningful variance associated with childhood maltreatment (CM) and diagnosis being removed by ComBat-GAM due to conflation with site effects (5). We therefore opted to report the non-harmonized results in the main manuscript.

Model robustness was confirmed through sensitivity analyses across varying age groups and longitudinal assessments over a two-year period. CentileBrain normative models demonstrated stable and high accuracy, plateauing at a sample size of approximately 3,000 participants. The finalized models are publicly accessible through the CentileBrain platform: <https://centilebrain.org/>

**Methods S4. Sex and age differences in associations between FreeSurfer metrics and CTQ**

To statistically compare the strength of associations between sexes and age cohorts, we employed Fisher's exact transformation tests. This method allowed us to determine whether the standardized regression coefficients (β) from our general linear models differed significantly between males and females (within the same age cohort) and between young adults and older adults (within the same sex). For each comparison, we computed a two-tailed *p*-value testing the null hypothesis that the effect sizes were equal between groups. We also determined which group demonstrated the stronger effect (based on the absolute magnitude of β values) for each brain region where significant associations were observed. These comparisons were performed separately for each morphometric phenotype (subcortical volume, cortical thickness, and surface area) and type of CM (abuse and neglect).

**Methods S5. Post-hoc analysis testing effect of diagnosis**

To account for potential confounding effects of psychiatric diagnoses, we conducted analyses adjusting for diagnostic status (“patients” versus “healthy controls”) as a covariate in our general linear models examining associations between CTQ scores and brain deviation scores. We grouped individuals into “patients” (including those diagnosed with MDD, PTSD, or comorbid presentations) versus “healthy controls” rather than separating MDD and PTSD. This approach was chosen because recruitment across study sites varied—with some sites enrolling predominantly MDD patients (who may also exhibit PTSD symptoms) and others enrolling PTSD patients (with potential overlap in depressive symptoms), as well as controls from PTSD sites who may have clinical-level MDD symptoms. Consequently, the binary grouping of patients versus controls serves as a rough proxy for diagnostic status, though these findings should be interpreted with caution given the inherent heterogeneity in determining diagnoses across sites.

**Methods S6. Testing site-specific effects using leave-one-out cross-validation**

Given varying sample sizes and characteristics across sites (see Table S2), we performed leave-one-out cross-validation to assess site-specific influences. In this procedure, we sequentially excluded one site at a time and re-ran our general linear models, thereby generating a set of *p-*values (one per site omitted). We then calculated the range of these *p*-values to evaluate the stability of our findings. The idea was that if the removal of a particular site led to a marked change in the significance of the association—for example, a shift from a statistically significant to a non-significant result—this would indicate disproportionate influence. Although we did not define a formal numerical threshold for “disproportionate” influence, we interpreted substantial variability in the *p*-value range (e.g., a change that effectively crossed the conventional significance boundary) as evidence of a site-specific effect. With this analysis, the *p*-values remained relatively consistent across iterations, suggesting that no single site had a disproportionate impact on the observed associations.

**Methods S7. Results with ComBat-GAM for site harmonization**

Given the multi-site nature of our dataset, comprising diverse scanners, demographic distributions, and clinical characteristics across 25 independent cohorts, site harmonization was performed using ComBat-GAM (generalized additive model) (5). ComBat-GAM adjusts for systematic scanner-related biases while preserving meaningful biological variance through an empirical Bayes framework that estimates and removes unwanted site effects from morphometric measures (e.g., cortical thickness, surface area, subcortical volumes). This enhances comparability across sites.

However, it is important to highlight that the application of ComBat-GAM is not ideal for our dataset due to the absence of psychiatrically healthy controls in some sites, potentially affecting the accuracy and generalizability of harmonization outcomes. Despite these limitations, ComBat-GAM harmonization was included in supplementary analyses to provide comprehensive and transparent evaluation. The harmonization was applied independently to each morphometric measure (Table S7). Nonetheless, primary analyses utilized non-harmonized data to preserve potentially meaningful variance linked to clinical characteristics and diagnostic status.

**Methods S8. Power analysis for pediatric cohort**

Given the smaller sample size of the pediatric cohort compared to those found in the young adult and older adult cohorts, we conducted a bootstrap sampling analysis to determine if the null findings in the pediatric cohort was possibly attributed to limited statistical power rather than true developmental differences. For each significant association identified in the young adult and older adult cohorts across all morphometric phenotypes, we randomly down-sampled the datasets to match the pediatric sample size (maintaining the same sex distribution) and performed the same statistical analyses across 500 bootstrap iterations. Statistical power was calculated as the proportion of bootstrap iterations yielding significant results (mean *p*<0.05), with a threshold of 0.8 (80%) considered adequate power.

**Supplemental Results**

**Figure S1. Geographical distribution of participating research sites**. Colors indicate the ENIGMA work group that the sites in each country contributed to: sites that contributed data to the ENIGMA Major Depressive Disorder Work Group (MDD, blue), ENIGMA Posttraumatic Stress Disorder Work Group (PTSD, yellow), or both ENIGMA Work Groups (green). Note that each country may have multiple research sites contributing to these ENIGMA Work Groups.

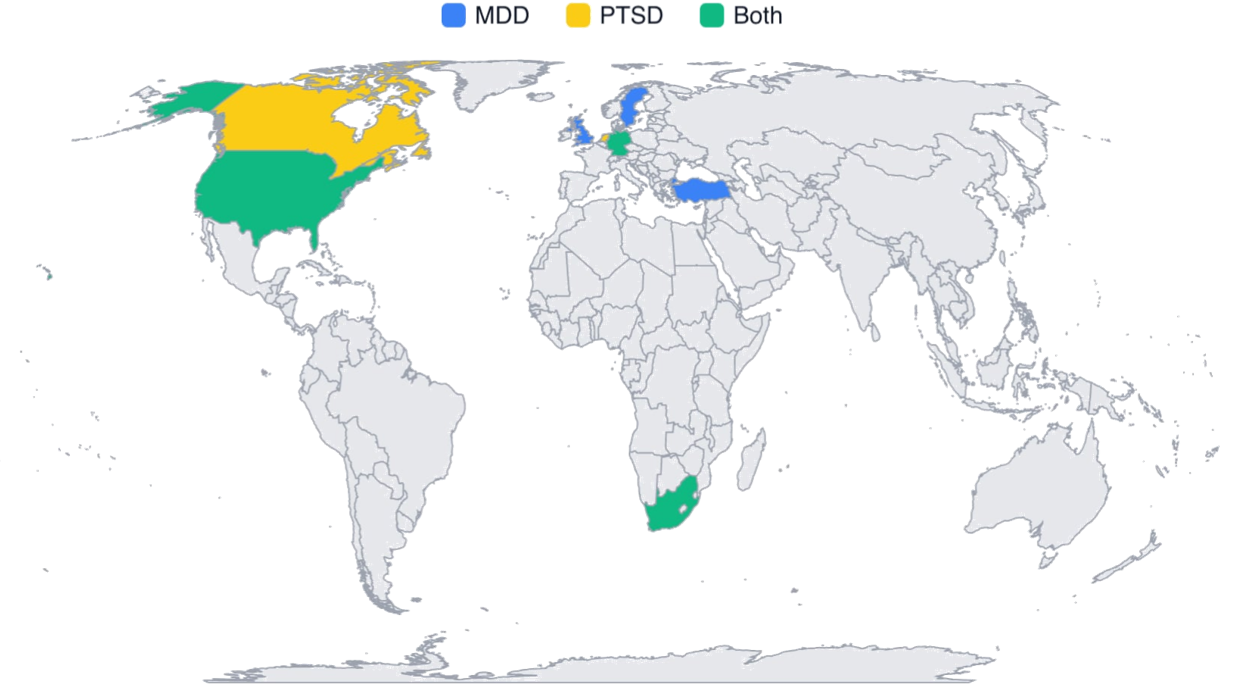

**Table S1A.** **ENIGMA site**s s**ample characteristics and demographics**

| **Site** | **Work Group** | **Age range** | **Mean age (SD)** | **Female %** | **MDD Sample Size** | **PTSD Sample Size** | **Controls Sample Size** |
| --- | --- | --- | --- | --- | --- | --- | --- |
| AFFDIS | MDD | 19.0-61.0 | 35.73 (14.35) | 46.90% | 29 | 0 | 20 |
| BRDECC | MDD | 33.0-62.0 | 51.57 (6.58) | 64.30% | 9 | 0 | 19 |
| CSAN | MDD | 18.0-65.0 | 34.70 (12.94) | 67.90% | 60 | 0 | 49 |
| FOR2107 | MDD | 18.0-65.0 | 34.47 (13.04) | 63.40% | 492 | 0 | 635 |
| Houston-Young | MDD | 10.0-17.0 | 12.86 (2.41) | 21.40% | 0 | 0 | 14 |
| INTRUST | PTSD | 18.0-69.0 | 35.70 (12.40) | 39.90% | 0 | 93 | 253 |
| Munster | MDD | 16.0-63.0 | 36.75 (11.88) | 55.60% | 182 | 0 | 544 |
| SoCAT | MDD | 18.0-26.0 | 23.15 (1.98) | 100.00% | 20 | 0 | 27 |
| Stanford-DTI | MDD | 18.9-52.1 | 32.78 (9.86) | 100.00% | 14 | 0 | 17 |
| Stanford-TIGER | MDD | 13.0-18.0 | 15.73 (1.34) | 64.70% | 40 | 0 | 11 |
| UCSF | MDD | 13.0-18.0 | 15.35 (1.43) | 50.80% | 26 | 0 | 39 |
| Cape Town | PTSD | 17.0-43.0 | 26.39 (6.46) | 100.00% | 0 | 1 | 154 |
| Columbia | PTSD | 18.0-60.0 | 35.63 (12.45) | 44.00% | 0 | 25 | 50 |
| Duke | PTSD | 24.0-67.0 | 40.96 (9.61) | 26.80% | 0 | 19 | 37 |
| Emory | PTSD | 22.0-62.0 | 40.85 (12.10) | 100.00% | 0 | 14 | 46 |
| Groningen | PTSD | 23.0-58.0 | 38.22 (9.68) | 100.00% | 0 | 36 | 0 |
| Mclean | PTSD | 18.0-62.0 | 33.59 (11.97) | 100.00% | 0 | 43 | 27 |
| Milwaukee | PTSD | 18.3-57.9 | 32.51 (10.38) | 49.30% | 0 | 21 | 54 |
| Ontario | PTSD | 18.0-60.0 | 35.55 (12.37) | 70.70% | 0 | 52 | 40 |
| Stanford | PTSD | 18.0-61.0 | 34.13 (11.08) | 45.20% | 0 | 67 | 68 |
| UW_Cisler | PTSD | 20.0-50.0 | 32.67 (8.11) | 100.00% | 0 | 65 | 10 |
| UWash | PTSD | 8.1-17.3 | 12.79 (2.64) | 49.30% | 0 | 32 | 114 |
| Vanderbilt | PTSD | 23.0-40.0 | 31.34 (4.63) | 18.00% | 0 | 15 | 35 |
| Waco_VA | PTSD | 26.0-56.0 | 39.77 (11.05) | 11.50% | 0 | 13 | 13 |
| Yale | PTSD | 20.0-52.0 | 30.19 (8.03) | 16.40% | 0 | 21 | 46 |

**Table S2.** **ENIGMA sites scan image acquisition and processing**

| **Site** | **Country** | **Scanner type** | **Sequence T1** | **FreeSurfer version** | **Slice orientation** | **Operating system** |
| --- | --- | --- | --- | --- | --- | --- |
| AFFDIS | Germany | 3T Siemens Magnetom TrioTim | 3D T1: 176 slices, TR = 2250 ms, TE = 3.26 ms, FOV 256, voxel size 1 x 1 x 1mm | 5.3 | Sagittal | Linux CentOS |
| BRDECC | UK | 1.5T GE Signa HDx | ADNI-1 MPRAGE pulse sequence | 5.3 | Sagittal | Linux-centos4_x86_64 |
| CSAN | Sweden | 3T Siemens MAGNETOM PRISMA | 3D T1: MPRAGE, TR = 2300 ms, TE = 2.34 ms, FOV 250 x 250 mm, voxel size = 0.9 x 0.868 x 0.868 mm, flip angle = 8° | 7.2 | Sagittal | Ubuntu |
| FOR2107 | Germany | 3T Siemens Magnetom TiroTim syngo MR B17 | 3D T1: MPRAGE, 176 slices, Slice Gap = 0.5 mm, voxel size 1 x 1 x 1mm, TI = 900 ms, TE = 2.26 ms, TR = 1900 ms, flip angle = 9° | 5.3 | Sagittal | Red Hat Enterprise Linux Server release 5.11 (Tikanga) |
| Houston-Young | US | 1.5 T Philips Medical Systems Gyroscan Intera; 3T Siemens Allegra | 1.5T: Fast field echo sequence, TR = 24 ms, TE = 4.99 ms, flip angle = 40°, slice thickness = 1 mm, FOV = 256 × 256, 150 slices.  3T: MPRAGE, TR = 1750 ms, TE = 4.39 ms, flip angle = 8°, slice thickness = 1 mm, FOV = 208 × 256, 160 slices. | 5.3 | Sagittal; Transverse | Fedora 19 |
| INTRUST | US | GE; Siemens; Philips | Multiple T1 protocols: SPGR-BRAVO/MPRAGE/ T1W_3D_TFESE NSE, voxel size = 1 x 1 x1, FOV = 256 x 256 mm, TR = 9150/2530/7600 ms, TE = 3.7/3.32/3.5 ms, flip angle = 10°/7°, slice thickness = 1mm | 5.3 | Sagittal | Linux |
| Munster | Germany | 3T Philips Gyroscan Intera | 3D T1: fast gradient echo sequence (turbo field echo), TR = 7.4 ms, TE = 3.4 ms, flip angle = 9°, 2 signal averages, inversion prepulse every 814.5 ms, FOV = 256 x 204 x 160 mm, voxel size = 0.5 x 0.5 x 0.5 mm | 5.3 | Sagittal | Red Hat Enterprise Linux Server release 5.11 (Tikanga) |
| SoCAT | Turkey | 3T Siemens Verio,Numaris/4;Syngo MR B17 | 3D T1: MPRAGE; TR=1900 msec; TE=3.4 msec; Flip angle=15°; Voxel size 1 mm x 1 mm x 1 mm | 7.1 | Axial | freesurfer-linux-centos7_x86_64-7.1.1 |
| Stanford-DTI | US | 3.0T GE Discovery MR750 | T1: SPGR, 186 slices, resolution = 0.9 mm isotropic, flip angle = 12°, TR = 6,240 ms; TE = 2.34 ms | 5.3 | Sagittal | Centos6_x86_64, Linux-based HPC |
| Stanford-TIGER | US | 3T GE MR750 | TR/TE/TI=8.2/3.2/600 ms; flip angle = 12°, 156 slices; FOV=256 x 256 mm, isotropic voxel = 1 mm, total scan time = 3:40 | 6 | Axial | Linux |
| UCSF | US | 3T GE Discovery MR750 | T1: SPGR, TR = 8.1 ms; TE = 3.17 ms; TI = 450 ms; flip angle = 12°; 256 x 256 matrix, FOV = 250 x 250 mm, 168 slices; slice thickness = 1 mm, in-plane resolution = 0.98 x 0.98 mm | 5.3 | Sagittal | Linux-centos6_x86_64-stable-pub-v5.3.0 |
| Cape Town | South Africa | Siemens Skyra | 3T T1: MPRAGE, voxel size = 1 x 1 x 1.5 mm, FOV = 256 x 256 mm, TR = 2530 ms, TE = 1.69/3.55/5.41/7.27 ms, flip angel = 7° | 7.3.2 | Sagittal | Linux |
| Columbia | US | GE Signa Excite | 1.5T T1: MPRAGE, voxel size = 1 x 1 x 1.3 mm, FOV = 256 x 256 mm, TR = 7250 ms, TE = 3 ms, flip angle = 7° | 7.3.2 | Axial | Linux |
| Duke | US | GE Discovery MR750; GE LX Nvi; Philips Ingenia; GE Signa EXCITE | 3T/4T T1: FSPGR/FSPGR BRAVO/Spin-echo co-planar/  3D TFE SENSE, voxel size = 1 x 1 x 1/0.9375 x 0.9375 x 1/1 x 1 x 1.9 mm, FOV = 240 x 240/256 x 256 mm, TR = 3.22/7.84/8.16/12/8.148/8.208 ms, TE = 8.148/2.98/3.18/5.4/3.728/3.22 ms, flip angle = 12°/20°/8° | 7.3.2 | Axial | Linux |
| Emory | US | Siemens TIM Trio | 3T T1: MPRAGE, voxel size = 1 x 1 x 1 mm, FOV = 224 x 256 mm, TR = 2600 ms, TE = 3.02 ms, flip angle = 8° | 7.3.2 | Axial | Linux |
| Groningen | Netherlands | Siemens TIM Trio | 3T T1: MPRAGE, voxel size = 1 x 1 x 1 mm, FOV = 256 x 256 mm, TR = 1900 ms, TE = 2.52 ms, flip angle = 9° | 7.3.2 | Sagittal | Linux |
| Mclean | US | Siemens TIM Trio | 3T T1: MPRAGE, voxel size = 1.3 x 1.3 x 1.3 mm, FOV = 256 x 128 mm, TR = 2530 ms, TE = 3.31 ms, flip angle = 7° | 7.3.2 | Sagittal | Linux |
| Milwaukee | US | GE Discovery MR750 | 3T T1: SPGR, voxel size = 1 x 0.9375 x 0.9375 mm, FOV = 240 x 240 mm, TR = 9800 ms, TE = 4.6 ms, flip angle = 8° | 7.3.2 | Sagittal | Linux |
| Ontario | Canada | Siemens Biograph mMR | 3T T1: MPRAGE, voxel size = 1 x 1 x 1 mm, FOV = 256 x 240 x 192 mm, TR = 2300 ms, TE = 2.98 ms, flip angle = 9° | 7.3.2 | Axial | Linux |
| Stanford | US | GE Discovery MR750 | 3T T1: SPGR, voxel size = 1.5 x 0.9 x 1.1 mm, FOV = 220 x 220 / 240 x 240 mm, TR = 8000/8600 ms, TE = 3.6/3.4 ms, flip angle = 15° | 7.3.2 | Coronal | Linux |
| UW_Cisler | US | Philips Achieva X-Series;  GE Discovery MR750 | 3T T1: MPRAGE, voxel size = 1 x 1 x 1 mm, FOV = 256 x 256 mm, TR = 8200 ms, TE = 3.2 ms, flip angle = 12° | 7.3.2 | Sagittal; Axial | Linux |
| UWash | US | Philips Achieva | 3T T1: MPRAGE, voxel size = 1 x 1 x 1 mm, FOV = 256 x 256 mm, TR = 2530 ms, TE = 3.5 ms, flip angle = 7° | 7.3.2 | Transversal | Linux |
| Vanderbilt | US | Philips Intera | 3T T1: voxel size = 0.8 x 0.8 x 0.9 mm, FOV = 256 x 256 mm, TR = 9000 ms, TE = 4.6 ms, flip angle = 9° | 7.3.2 | Sagittal | Linux |
| Waco_VA | US | Philips Achieva | 3T T1: MPRAGE, voxel size = 0.9 x 0.9 x 0.9 mm, FOV = 256 x 256 mm, TR = 7256 ms, TE = 2.77 ms, flip angle = 12° | 7.3.2 | Sagittal | Linux |
| Yale | US | Siemens TIM Trio | 3T T1: MPRAGE, voxel size = 1 x 1 x 1 mm, FOV = 256 x 256 mm, TR = 2500 ms, TE = 2.77 ms, flip angle = 7° | 7.3.2 | Sagittal | Linux |

**Note:** FOV=field of view, TR=repetition time, TE=echo time, T=Tesla, FSPGR=fast spoiled gradient echo, BRAVO = brain volume, TFE=turbo field echo, SENSE=sensitivity encoding, MEMPRAGE=multi-echo MPRAGE, MPRAGE=magnetization prepared rapid gradient echo, TFL=tensor fascia lata, SPGR=spoiled gradient echo, EPI=echo planar.

**Table S3. ENIGMA istrument for clinical diagnosis and local study eligibility criteria.**

**Table S3A. ENIGMA - MDD Working Group instrument for clinical diagnosis and study eligibility criteria by site**.

| **Site** | **Diagnostic instrument** | **Inclusion criteria** | **Exclusion criteria** |
| --- | --- | --- | --- |
| AFFDIS | ICD-10, DSM-IV criteria | MDD subjects were currently depressed and in day program or inpatient | All subjects exclusion criteria: current or history of neurological disorder or brain injury, current substance abuse or dependence (not including nicotine), pregnancy, MRI contraindications, inability to give consent. MDD specific: comorbid psychiatric diagnosis. Healthy control specific: current or history of psychiatric diagnosis. |
| BRDECC | SCAN | Community based or outpatients with less than two depressive episodes of at least moderate severity. Did not meet DSM-IV diagnostic criteria for recurrent major depressive disorder.  Control group participants were clinically interviewed to ensure they had never experienced depressive symptoms. | Contraindications to MRI, diagnosis of neurological disorder, head injury leading to loss of consciousness or conditions known to affect brain structure or function (including alcohol or substance misuse), if they or a first-degree relative had ever fulfilled criteria for mania, hypomania, schizophrenia or mood-incongruent psychosis. |
| CSAN | MINI | MDD subjects were currently depressed meeting MINI criteria for depression; comorbid anxiety disorders are allowed; mood-congruent psychotic symptoms allowed. | Current MDD: a current DSM-5 diagnosis of substance use disorder, except nicotine; a psychotic disorder, except depression with mood-congruent psychotic features; new antidepressant medication during the month before study participation (two months for fluoxetine); change of the dose of psychotropic medications over the last month (antidepressant and antipsychotic medication) or the last two months (mood stabilizers and anticonvulsants). |
| FOR2107 | SCID | Participants recruited by means of public advertisement and from the inpatient services. Inclusion criteria: age 18-65 years; patients were diagnosed with major depressive disorder by SCID-Interview, currently depressed or remitted. | Any MRI contraindications; any neurological abnormalities. Exclusion criteria controls: any current or former psychiatric disorder; Exclusion criteria patients: substance dependence or current benzodiazepine treatment (wash out of at least three half-lives before study participation) |
| Houston-Young | DSM-IV criteria | Outpatients with MDD diagnosis according to DSM-IV | MDD subjects: head trauma with residual effects, neurological disorders, uncontrolled major medical conditions based on patient self reports and current drug abuse. In addition, Healthy controls were excluded if they had a history of any Axis I disorder or had a first-degree relative with any Axis I disorder. |
| Munster | SCID | Participants recruited by means of public advertisement and from the inpatient services. Inclusion criteria: age 16-65 years; patients were diagnosed with MDD by SCID-Interview | MDD subjects: presence of bipolar disorder, schizoaffective disorders and schizophrenia; substance-related disorders or current benzodiazepine treatment (wash out of at least three half-lives before study participation), and former electroconvulsive therapy. Control subjects: any current or former psychiatric disorder. Both groups: any neurological abnormalities, MRI contra-indications |
| SoCAT | SCID | DSM IV diagnosis for MDD patients. Age: 18-65. Right-handed, currently depressed or remitted; Control subjects: no any history of psychiatric disorder | Exclusion criteria 1) History of significant head injury 2) Neurological diseases such as epilepsy, cerebrovascular accident 3) Other diagnoses on Axis I disorders4) |
| Stanford-DTI | SCID | Community-based DSM-diagnosed sample | MDD subjects: presence of axis-I disorders other than MDD, anxiety and eating disorders . Control subjects: control individuals did not meet diagnostic criteria for any current psychiatric. Both groups: alcohol / substance abuse or dependence within six months prior to MRI scanning, history of head trauma with loss of consciousness > 5 min, aneurysm, or any neurological or metabolic disorders that require ongoing medication or that may affect the central nervous system (including thyroid disease, diabetes, epilepsy or other seizures, or multiple sclerosis), MRI contraindications, or bad MRI data (e.g., extreme movement). |
| Stanford-TIGER | KSADS | Community-based DSM-diagnosed sample | All subjects: Exclusion criteria were premenarchal status (for females), history of concussion within the past 6 weeks or history of any lifetime concussion with loss of consciousness, contraindications to MRI scanning (e.g. braces, metal implants, or claustrophobia), serious neurological or intellectual disorders that could interfere with the participant's ability to complete study components. MDD subjects: meeting lifetime or current DSM-IV criteria for any Bipolar Disorder, Psychosis, or Alcohol Dependence, or DSM-5 criteria for Moderate Substance Use Disorder with substance-specific threshold for withdrawal. CTL subjects: any current or past DSM-IV Axis I Disorder and first-degree relative with confirmed or suspected history of depression, mania, psychosis, or substance dependence. |
| UCSF | KSADS for MDD, DISC for controls | Outpatient/community-based sample with DSM diagnosis, mostly antidepressant-naive and approximately 60% of MDD have comorbid anxiety disorders | Exclusion criteria for all participants included: 1) use of pharmacotherapeutics for treating psychiatric conditions within the past 6 months, 2) misuse of drugs within two months prior to MRI scanning; 3) two or more alcoholic drinks per week within the previous month (as assessed by the CDDR); 4) a full scale IQ score of less than 75; 5) contraindications for MRI including ferromagnetic implants and claustrophobia; 6) pregnancy or the possibility of pregnancy; 7) left-handedness; 8) prepubertal status (as assessed as Tanner stages of 1 or 2) (Tanner, 1962); 9) inability to understand and comply with procedures; 10) neurological disorder (including meningitis, migraine, or HIV); 11) head trauma; 12) learning disability; 13) serious health problems; and 14) complicated or premature birth (i.e., birth before 33 weeks of gestation). The MDD group was subject to the additional exclusion criterion of a primary psychiatric diagnosis other than MDD. The control group was subject to the additional exclusion criteria of: 1) history of mood or psychotic disorders in a first- or second-degree relative; and 2) current or lifetime DSM-IV-TR Axis I psychiatric disorder. |

**Note:** MDD = Major Depressive Disorder, SCAN = Schedules for Clinical Assessment in Neuropsychiatry, MINI = Mini International Neuropsychiatric Interview, SCID = Structured Clinical Interview for DSM, DSM-IV = Diagnostic and Statistical Manual of Mental Disorders, Fourth Edition, DSM-IV-TR = Diagnostic and Statistical Manual of Mental Disorders, Fourth Edition, Text Revision, KSADS = Kiddie Schedule for Affective Disorders and Schizophrenia, CDDR = Customary Drinking and Drug Use Record. DISC = Diagnostic Interview Schedule for Children, CTL = Controls.

**Table S3B. ENIGMA - PTSD Working Group instrument for clinical diagnosis and study eligibility criteria by site.**

| **Site** | **Diagnostic instrument** | **Inclusion criteria** | **Exclusion criteria** |
| --- | --- | --- | --- |
| INTRUST | DSM-5 Criteria | Male and female civilians or veterans. | Healthy controls: any trauma exposure. All: contraindications to MRI |
| Cape Town | DSM-5 Criteria | Women 18-65 years of age who speak English, Afrikaans, or Xhosa | Intellectual disability; critical medical condition; current psychotic episode/disorder; contraindications for MRI |
| Columbia | DSM-5 Criteria | All: Males or females 18-60 years of age able to give consent, fluent in English  PTSD: Experience of a traumatic event or events during lifetime; current DSM-5 Criterion A for PTSD | All: history of psychosis, bipolar disorder, or dementia; significant depression (HAM-D>25); suicidality; recent substance/alcohol dependence (past 6 months) or abuse (past 2 months); psychotropic medication usage within past 4 weeks (e.g., antipsychotics, antidepressants, mood stabilizers, or stimulants); triptan anti-migraine medications; β-blockers; pregnancy; MRI contraindications; serious and/or unstable/untreated medical illness (e.g., stroke, brain tumor, demyelinating disease)  Controls: Current or lifetime history of major psychiatric diagnosis, e.g., major depressive disorder, psychotic disorder, bipolar disorder, obsessive compulsive disorder (OCD), PTSD, panic disorder, agoraphobia, eating disorder or alcohol/substance use disorder; history of DSM-5 criterion A1 trauma exposure; HAM-D>7 |
| Duke | DSM-5 Criteria | Veterans 18-65 years of age, fluent in English, free of implanted metal objects or metal shards in eyes | Axis I psychiatric conditions other than PTSD or MDD; current substance abuse or lifetime substance dependence (other than nicotine); high risk for suicide, claustrophobial; neurological disorders; learning disability or developmental delay; major medical conditions |
| Emory | DSM-5 Criteria | Individuals 18-65 years of age who speak English and have endorsed at least 1 criterion A trauma | Current psychotic symptoms or bipolar disorder; current substance or alcohol dependence; history of head trauma; psychoactive medication usage; current illegal drug use (verified with urine drug screen within 24 hours of scan) |
| Groningen | DSM-5 Criteria | Civilian women 20-60 years of age with a current PTSD diagnosis, sufficient proficiency in German, MRI compatible | Neurologic disorders; history of substance abuse or dependence (past 6 months); history of head injury; cerebral incidental findings verified by a neuroradiologist after the MR scan; usage of benzodiazepines, tricyclic antidepressants, or anticonvulsants; primary borderline personality disorder; other current diagnosis of Axis I disorder |
| Mclean | DSM-5 Criteria | Women 18-60 years of age with a history of childhood maltreatment who speak English; must have legal and mental competency, Normal or Corrected Vision. | Delirium secondary to medical illness; History of neurological conditions that may cause significant psychiatric symptomatology (e.g., dementia); Any contraindication to MR scans, including claustrophobia, pregnancy, metal implants, etc.; Current alcohol or substance use disorder (within the last month); A history of schizophrenia or other psychotic disorder; History of head injury or loss of consciousness for longer than 5 min (including concussion); pregnancy |
| Milwaukee | DSM-5 Criteria | Civilians aged 18-60 years; exposure to DSM-5 A1 criterion trauma; high risk for PTSD (score ≥3 OR item 2 rated ≥3 on Predicting PTSD Questionnaire, Rothbaum et al., 2014); English speaking; ability to schedule baseline study visit within 30 days of traumatic injury | Glasgow Coma Scale score ≤ 13 (i.e., moderate to severe traumatic brain injury); on police hold; contraindication to MRI; pregnancy (or planned pregnancy within 6 months); intentional self-inflicted injury; severe vision or hearing impairment; history of psychotic or manic symptoms, or neurologic condition (e.g., seizures, spinal cord injury); currently on antipsychotic medication; clear evidence of substance use disorder |
| Ontario | DSM-5 Criteria | Primary diagnosis of PTSD for patients | Incompatibilities with scanning conditions, previous neurologic and development illness, comorbid schizophrenia or bipolar disorder, alcohol or substance abuse, a history of head trauma, or pregnancy during scan. participants were excluded if they had implants or metal that do not comply with 3T fMRI safety standards for research, a history of head injury with a loss of consciousness, significant untreated medical illness, a history of neurological disorders, history of any pervasive developmental disorders, pregnancy, and current use of any psychotropic medication within one month prior to study. PTSD individuals were further excluded if they reported a history of bipolar disorder, schizophrenia, or substance-use disorder prior to participation of the study |
| Stanford | DSM-5 Criteria | All: Community-dwelling adults 18-60 years of age who are fluent in English  Patients: Individuals experiencing chronic (>3 months) moderate to severe anxiety or depression (indicated by a score >10 on the PHQ-9 [excluding suicide item] OR a score >10 on the GAD-7 scale) - must express interest in seeking treatment for psychiatric symptoms.  Controls: PHQ-9 and GAD ≤ 4. | All: contraindications to MRI or TMS; current participation in psychiatric treatment; history of neurological disorder, brain surgery, electroconvulsive or radiation treatment, brain hemorrhage or tumor, stroke, epilepsy, hypo- or hyper-thyroidism; medication use that substantially reduces seizure threshold to TMS (e.g., olanzapine, chlorpromazine, lithium) and unwillingness or inability to safely withdraw at least two weeks prior to TMS appointment; medications that interfere with blood flow (e.g., opiates, antihypertensive medication); insufficiently controlled thyroid dysfunction; current substance dependence (within past 3 months); refusal to abstain from illicit drug use for study duration; refusal to abstain from alcohol within 24 hours of MRI scans; pregnancy; prior exposure to deep brain stimulation, rTMS, or tDCS therapies; significant traumatic brain injury (indicated by loss of consciousness, post-trauma amnesia, imaging findings, penetrating brain injury); history of psychotic or manic symptoms. |
| UW_Cisler | DSM-5 Criteria | Women aged 21-50 with and without history of interpersonal violence – must be fluent in English | History of psychosis, medication changes within past 4 weeks, cognitive impairment, current substance or alcohol use disorder |
| UWash | DSM-5 Criteria | 8-20 years old English speaking | Psychiatric medication use (excepting stimulant meds for ADHD, which were discontinued for the scan). MRI contra-indications including braces, or other metal in the body or claustrophobia. Active substance dependence, pervasive developmental disorder, active safety concerns. |
| Vanderbilt | DSM-5 Criteria | OEF/OIF/OND Veterans 18-50 years of age who are fluent in English | Psychoactive medication usage in past 6 weeks; participated in psychotherapy within the past month; current substance use disorder (>6 month remission); positive urine drug or alcohol breath screen on MRI study day; history of psychotic or bipolar disorder, traumatic brain injury, or significant medical (e.g., cancer, HIV) or neurological illness (e.g., stroke, brain tumor, multiple sclerosis, epilepsy); contraindication to MRI  Trauma-exposed controls: lifetime diagnosis of PTSD; symptoms of hypervigilance  Healthy controls: any trauma exposure |
| Waco_VA | DSM-5 Criteria | Veterans 18-60 years of age with a clinical diagnosis of TBI in their VA medical record | Diagnosis of schizophrenia, schizoaffective disorder, bipolar disorder type I, severe substance use disorder, or a high risk of suicide; seizure disorder, dementia; absence of qEEG more than 2SDs outside of population means of healthy age-matched controls; MRI contraindications |
| Yale | DSM-5 Criteria | Individuals 21-60 years of age; at least one deployment on combat tour | Diagnosis of bipolar disorder or psychotic disorder; current benzodiazepine use; a history of ADHD, learning disorder, moderate or severe TBI, brain tumor, epilepsy, or a neurological disorder; current inpatient status; MRI contraindication. |

**Notes:** The ENIGMA PTSD Work Group has administered clinical data harmonization and converted different clinical measures to DSM-5 criteria. For sites where we had item level responses for a PTSD measure these DSM-IV symptoms were categorized according to DSM-5 criteria. For sites with subscale level data using CAPS-IV, PCL-C, or PCL-M, cluster B was carried into analyses and cluster D was renamed as cluster E. subscale scores for clusters C and D were considered missing (6). MRI = Magnetic Resonance Imaging, PTSD = Posttraumatic Stress Disorder, DSM-5 = Diagnostic and Statistical Manual of Mental Disorders, Fifth Edition, HAM-D = Hamilton Rating Scale for Depression, MDD = Major Depressive Disorder, OCD = Obsessive-Compulsive Disorder, PHQ-9 = Patient Health Questionnaire-9, GAD-7 = Generalized Anxiety Disorder-7, TMS = Transcranial Magnetic Stimulation, rTMS = Repetitive Transcranial Magnetic Stimulation, tDCS = Transcranial Direct Current Stimulation, ADHD = Attention Deficit Hyperactivity Disorder, OEF = Operation Enduring Freedom, OIF = Operation Iraqi Freedom, OND = Operation New Dawn, TBI = Traumatic Brain Injury, VA = Veterans Affairs, qEEG = Quantitative Electroencephalogram, CAPS-IV = Clinician-Administered PTSD Scale for DSM-IV, PCL-C = PTSD Checklist – Civilian Version, PCL-M = PTSD Checklist – Military Version.

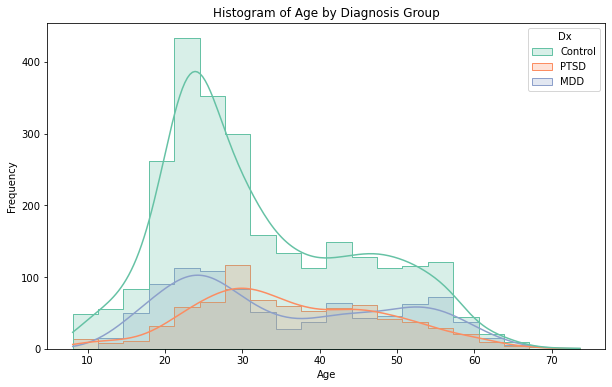
**Figure S2.** **Distribution of age by diagnosis.**

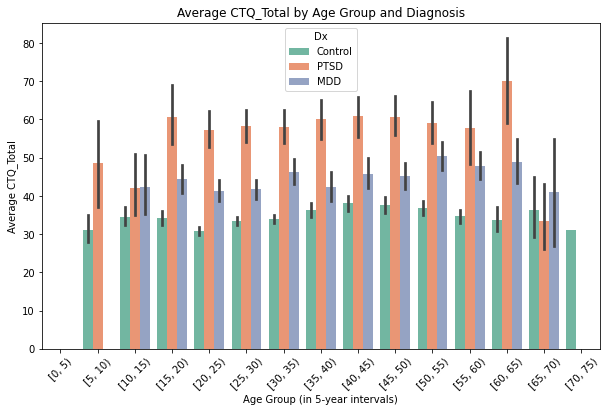
**Figure S3. Distribution of CTQ scores by age and diagnosis.**

**Table S4. Full results of normative deviations associated with CTQ.** Due to its size, this table is provided as a separate CSV file, accessible at https://github.com/haleyrwang/ENIGMA_CTQ/blob/main/Table_S4_CTQ_normative_deviations.csv.

**Table S5.** **Fisher's exact tests on normative deviations between age cohorts and between sex.** Due to its size, this table is provided as a separate CSV file, accessible at https://github.com/haleyrwang/ENIGMA_CTQ/blob/main/Table_S5_Fishers_comparisons.csv.

Fisher's r-to-z transformation tests revealed significant differences in the strength of associations between brain morphometric measures and CM exposure across demographic groups. Overall, young adults demonstrated stronger associations than older adults across all three structural phenotypes and both types of CM exposure, consistent with a sensitive period model of CM effects. Additionally, females generally showed stronger associations than males, particularly for surface area measures.

For subcortical volumes, comparisons across sexes revealed that most significant regions showed different effect strengths between males and females, with females consistently demonstrating stronger associations in both young adult and older adult cohorts for both abuse and neglect exposure. When comparing age cohorts, the young adult group consistently showed stronger associations than the older adult group for abuse- and neglect-related subcortical volume alterations.

For cortical thickness, sex differences followed a more complex pattern. In young adults, males showed stronger associations with childhood abuse, while in older adults, females exhibited stronger associations. For neglect, there were no significant sex differences in effect strengths in either age cohort. Across age groups, young adults demonstrated stronger associations between CM and cortical thickness compared to older adults for both abuse and neglect.

Surface area measures showed the most pronounced sex differences, particularly in young adults. For both abuse and neglect exposure, females demonstrated significantly stronger associations than males across a widespread set of cortical regions in the young adult cohort. This pattern persisted but was less extensive in the older adult cohort. Consistent with other morphometric measures, young adults showed stronger associations between CM and surface area compared to older adults across both sexes

**Table S6.** **Full results of normative deviation-CTQ association controlling for diagnosis.** Due to its size, this table is provided as a separate CSV file, accessible at https://github.com/haleyrwang/ENIGMA_CTQ/blob/main/Table_S6_CTQ_normative_deviations_dx_cov.csv.

To account for how associations between CM and structural brain deviation scores might differ based on clinical status, we conducted a variation of our main analysis (see Supplementary Method 5). Our analysis approach examined two complementary questions: (1) the effects of CM when adjusting for diagnostic status, and (2) the effects of diagnostic status when adjusting for CM. Specifically, we performed partial correlations between CM and structural brain deviation scores while controlling for diagnosis as a dichotomous variable (patient vs. control). We opted to group MDD and PTSD patients together to inform our transdiagnostic approach, and because the subject groups were not defined in the same way during recruitment across study sites.

When examining CM effects while adjusting for diagnosis, results were largely consistent with our primary findings across subcortical volume, cortical thickness, and surface area for the pediatric and young adult cohorts. While the associations between childhood abuse and brain deviation scores remained significant across young and older adult females even after accounting for diagnosis, the associations between childhood neglect and brain deviation scores that were unique to older adult females (i.e., those not also found in young adult females) - specifically, the associations with subcortical volumes and cortical thickness - were no longer significant after accounting for diagnosis.

When examining diagnosis effects while adjusting for CM, we found that diagnosis itself was associated with widespread cortical thickness deviations in pediatric and older adult female cohorts (but not male cohorts), particularly in bilateral frontal, temporal, and occipital regions. In our analyses relating CM to deviations in surface area, adjusting for diagnosis eliminated previously significant associations in young adult cohorts for both sexes; however, the null findings in the pediatric cohort and the significant results in older adult cohort remained unchanged. Overall, these sensitivity analyses on diagnostic effects demonstrates that most associations between CM and structural brain deviation scores are present even when accounting for psychiatric diagnosis, even though MDD and PTSD diagnoses (which could include treatment exposures) may be linked to additional brain morphometric deviations in early life that are independent of CM exposure.

**Table S7. Leave-one-out analysis results demonstrating the robustness of findings across sites.** Due to its size, this table is provided as a separate XLSX file, accessible at https://github.com/haleyrwang/ENIGMA_CTQ/blob/main/Table_S7_LOO_results.xlsx.

**Figures S4. Main results with ComBat-GAM harmonization.** CM +/- groups (i.e., Abuse +/-, Neglect +/-) were defined based on CTQ clinical cut-offs.

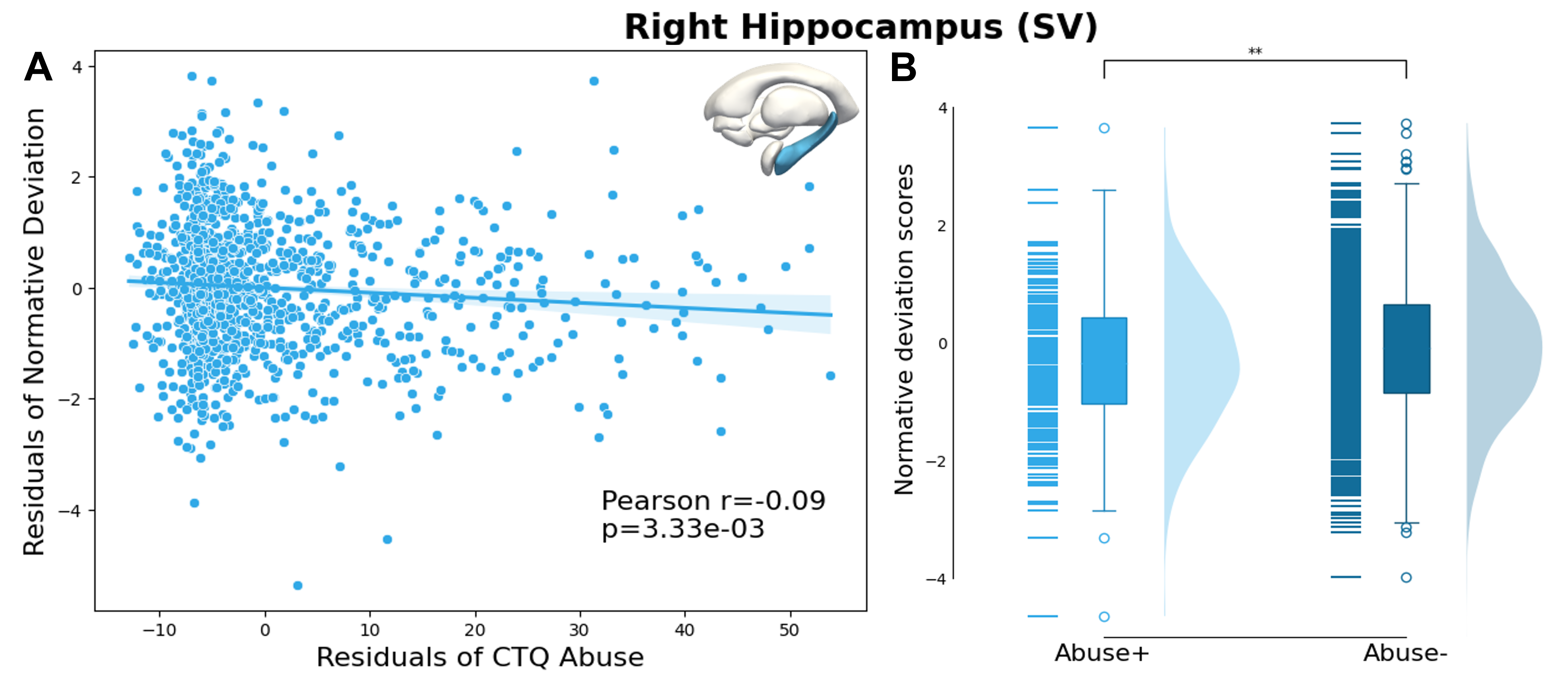

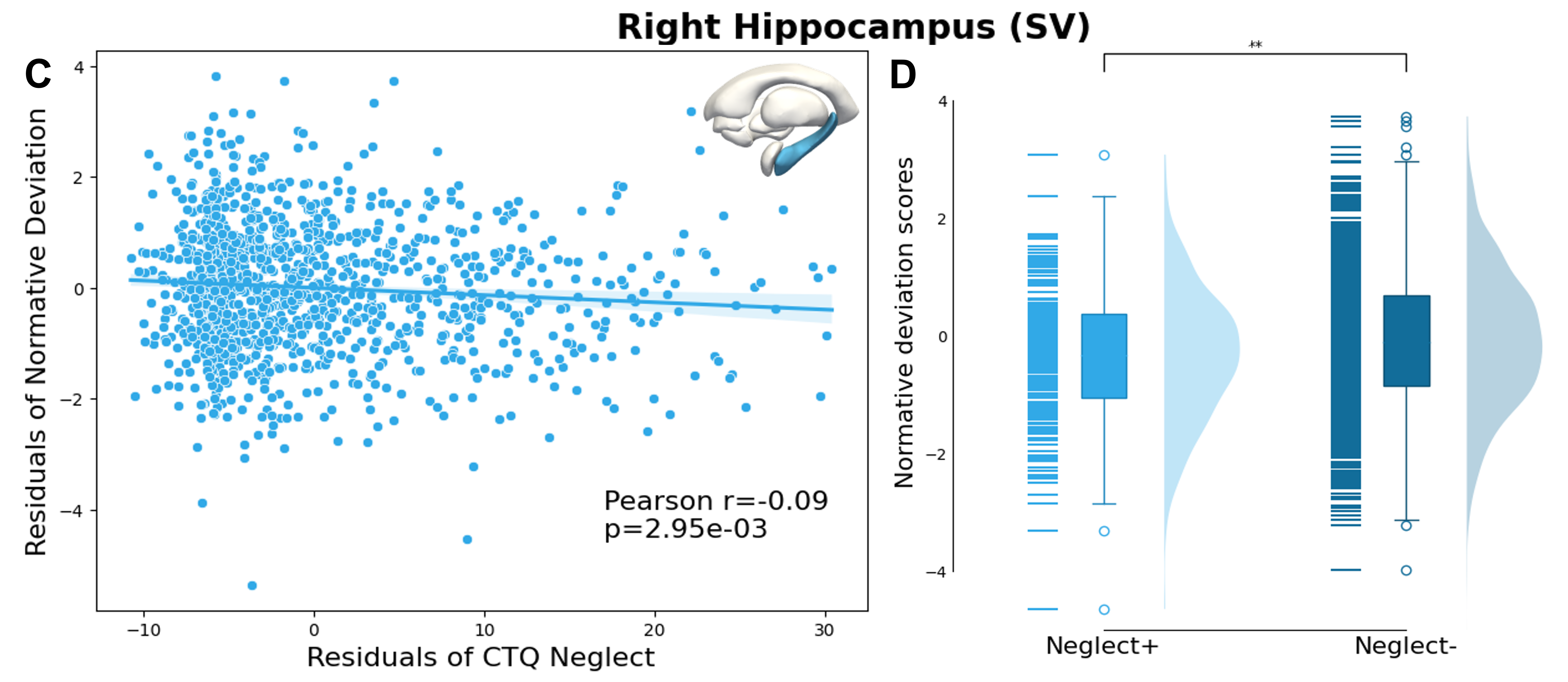

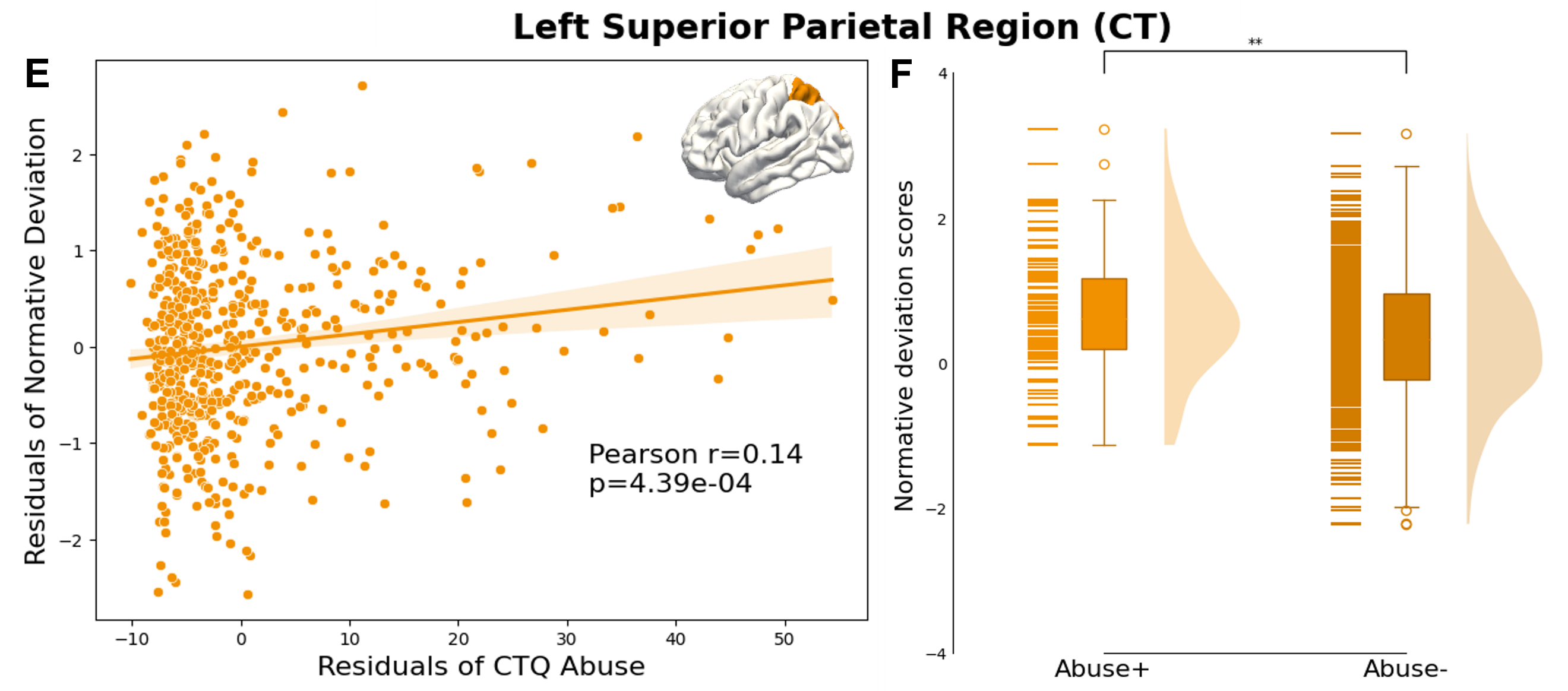

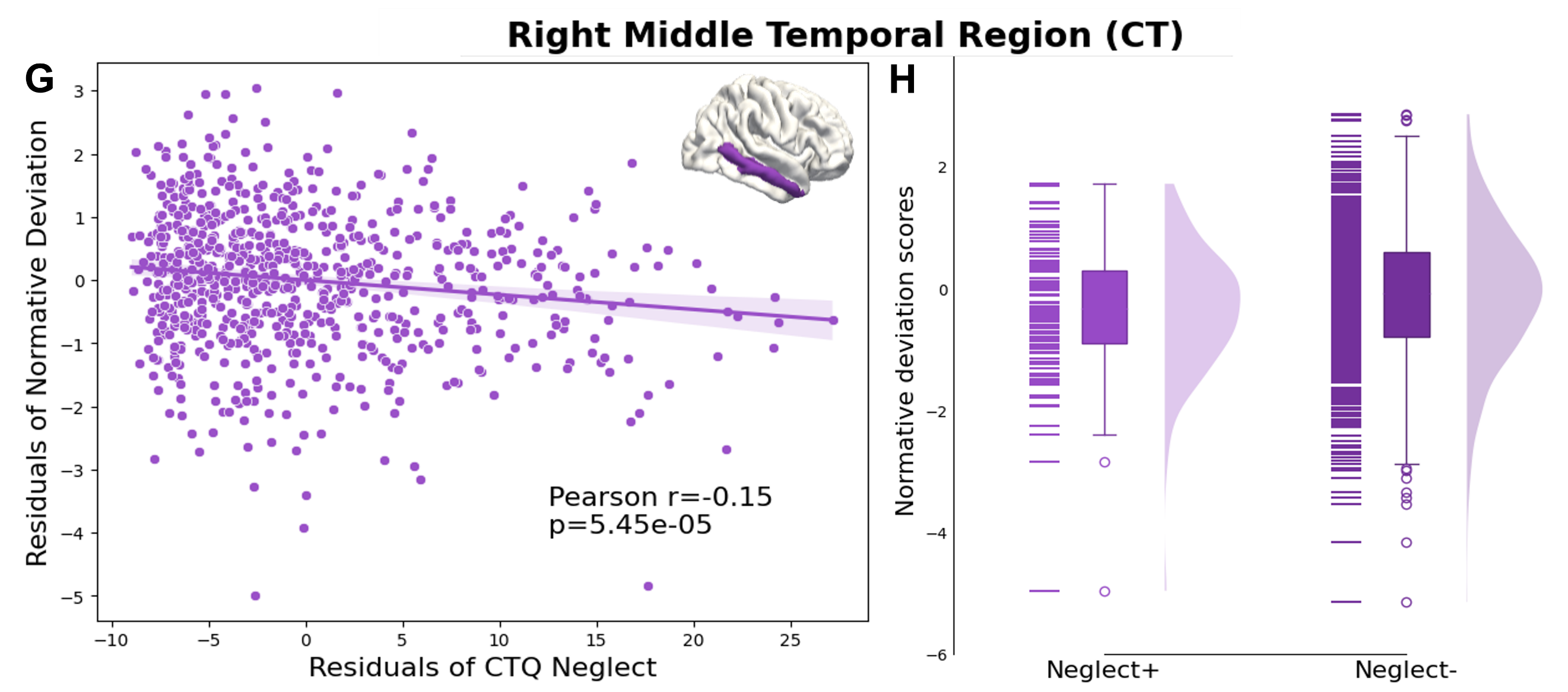

Among young adult females, both abuse (r=−.09, q<.05) and neglect (r=−.09, q<.05) were associated with decreased right hippocampal volume (Fig. S4A, S4C). Similarly, young adult females with moderate to severe abuse (t=-3.1) and neglect (t=-3.3) have significantly lower right hippocampal volumes compared to those with none to low trauma (Fig. S4B, S4D).

In older adult males, abuse correlated with increased cortical thickness in the left superior parietal region (r=.14, q<.05) (Fig. S4E). Older adult males with moderate to severe abuse have significantly thicker left superior parietal regions compared to those with none to low abuse (t=3.5) (Fig. S4F).

In young adult males, neglect correlated with thinner right middle temporal region (r=−.14, q<.05) (Fig. S4G). No other CM group difference on SV, CT, or SA was found (Fig. S4H).

**Result S1. Power analysis for pediatric cohort**

The bootstrap down-sampling analysis revealed that 3.4% of the significant associations observed in young adults and older adults cohorts would be reliably detected with the pediatric cohort sample size, significantly below the conventional power threshold of 80%. Among these few detectable associations were associations between childhood abuse and surface area in specific frontal regions (left frontal pole in young adult females and right medial orbitofrontal cortex in older adult females). The result from this analysis indicates that while null findings in children likely reflect limited statistical power rather than definitive age-related differences, the effects we identified in certain frontal regions are nonetheless robust enough to be detected with smaller samples
